# Cav3.1 is a leucine sensor in POMC neurons mediating appetite suppression and weight loss

**DOI:** 10.1101/2024.09.13.612843

**Authors:** Anthony H. Tsang, Nicholas Heeley, Constanza Alcaino, Eunsang Hwang, Brian Y. Lam, Taufiq Rahman, Tamana Darwish, Danae Nuzzaci, Richard G. Kay, Amar Sarkar, Ruiyan Wang, Nihal Basha, Austin Punnoose, Peter Kirwan, Marcella Ma, Giles S. Yeo, Florian T Merkle, Fiona M. Gribble, Frank Reimann, Kevin William, Clémence Blouet

**Affiliations:** Institute of Metabolic Science-Metabolic Research Laboratories, University of Cambridge, Cambridge, UK; Department of Internal Medicine, Center for Hypothalamic Research, Peter O’Donnell Jr. Brain Institute, University of Texas Southwestern Medical Center at Dallas, Dallas, Texas, USA; Department of Pharmacology, University of Cambridge, Cambridge, UK; Cambridge Stem Cell Institute, University of Cambridge, Cambridge, UK

**Author notes:** **Corresponding author**: Clemence Blouet. These authors contributed equally (Nicholas Heeley, Constanza Alcaino, Eunsang Hwang).

**Keywords:** Hypothalamus, Arcuate Nucleus, POMC neurons, Voltage-Gated Calcium Channel, Nutrient Sensing, Appetite, leucine, dietary proteins, Metabolic diseases, Obesity

## Abstract

Hypothalamic leucine sensing promotes satiety and weight loss but an understanding of how leucine regulates neuronal activity is lacking. Here we show that *Cacna1g*, encoding the T-type voltage-gated calcium channel Cav3.1, is enriched in hypothalamic leucine-sensing neurons and mediates leucine sensing. Pharmacological inhibition of Cav3.1 blunts leucine-induced activation of POMC neurons as well as the anorectic response to leucine in vivo. In addition, genetic deletion of *Cacna1g* in POMC neurons abolishes the appetite- and weight-suppressive effects of high-protein feeding. Mechanistically, we show that leucine binds to the voltage-sensing segment of Cav3.1, thereby reducing its threshold for voltage-dependent activation. Last, pharmacological activation of hypothalamic Cav3.1 promotes weight loss in diet-induced obese mice and potentiates the weight loss response to GLP-1 receptor agonism. These results reveal that Cav3.1 is a neuronal leucine sensor and a relevant weight loss target.

## Introduction

A rapid and efficient response to disturbances in nutrient levels is crucial for the survival of organisms from bacteria to humans. Cells have therefore evolved a host of molecular pathways that can sense nutrient concentrations and quickly regulate their activity and gene expression to adapt to changes in nutrient availability. In complex organisms, distributed populations of specialised nutrient-sensing cells continuously monitor nutrient availability and orchestrate a range of systemic metabolic responses to optimise inter-organ nutrient fluxes, allowing the maintenance of nutritional homeostasis. These include enteroendocrine cells in the gut, or endocrine cells of the pancreas. In the brain, monitoring of nutrient availability by specialised nutrient-sensing neurons provides an additional level of integrated regulation through the autonomic nervous system, and directly engages behavioural components recruited for nutritional homeostasis, i.e. hunger and satiety or macronutrient preference.

While the responses to shortage or excess in carbohydrate and fat are facilitated by the presence of storage compartments for these macronutrients in the liver and adipose tissue respectively, deviations in protein availability are optimally addressed through behavioural strategies in the absence of storage compartment for protein. However, how the brain senses protein availability to regulate appetite is poorly understood. It is well established that dietary proteins trigger the secretion of appetite-suppressing incretins during the digestion process ^1^, but postprandial excursions in circulating levels of specific amino acids also reach the brain to directly activate appetite-suppressing neurocircuits ^2,3^. These include the branched-chain amino acid Leucine, recognised as a signalling molecule reporting amino acid availability to regulate a variety of anabolic processes and produce satiety. However, the molecular identity of leucine-sensing neurons is poorly characterised. Leucine-sensing neurons represent a molecularly diverse population of neurons which includes pro-opiomelanocortin (POMC) neurons of the arcuate nucleus of the hypothalamus (ARH) ^4–6^. Leucine-induced mTOR signalling mediates leucine’s anorectic action ^4,7,8^, but POMC neurons rapidly respond to leucine independently of mTOR signalling through a mechanism that remains unclear ^6^.

To identify the molecular signature of hypothalamic leucine-sensing neurons, we employed an unbiased activity-dependent ribosome profiling technique, the PhosphoTRAP assay ^9^, which highlighted an enrichment of *Cacna1g* (calcium voltage-gated channel subunit alpha1 G) – encoding for a T-type voltage-gated calcium channel (VGCC) Cav3.1 – in leucine-sensing neurons. Further characterisation revealed a role for Cav3.1 as a direct molecular sensor for leucine in mediobasal hypothalamic (MBH) POMCs neurons and human-iPSC-induced POMC neurons, required for the appetite-suppressive action of high-protein feeding, one of the most successful and popular dietary strategy promoting satiety and weight loss ^10–12^. Capitalising on these findings, we found that pharmacological activation of MBH Cav3.1 promotes weight loss in diet-induced obese mice and potentiates the weight-loss effects of liraglutide, a GLP-1 receptor agonist. Together, our findings unravel an unexpected appetite-regulating central nutrient-sensing mechanism, leading us to devise a potential anti-obesity pharmacotherapeutic strategy.

## Results

### Molecular profiling of hypothalamic leucine-sensing neurons identifies *Cacna1g* as a top marker

To identify the molecular signature of leucine-activated neurons in the MBH, we adopted the PhosphoTRAP assay following parenchymal injection of leucine into the MBH as previously described (**Figure 1A**) ^5^. Neurons activated following MBH leucine injection (and expressing the neuronal activation marker cFos) co-express Ser240/244-phosphorylated ribosomal S6 protein (p-rpS6, **Figure S1A**), allowing selective immunoprecipitation-based capture and sequencing of polysomes from leucine-activated neurons. RNA sequencing of total MBH extracts (input sample) confirmed enrichment in ARH-specific genes, validating the neuroanatomical accuracy of the dissections (**Figure S1B**). Further, activity-dependent transcripts were significantly enriched in p-rpS6 immunoprecipitated samples (IP sample; **Figure S1C**), confirming enrichment in translating transcripts from activated neurons. Differential expression analysis between IP and input samples generated a list of enriched transcripts in leucine-activated neurons (**Figure 1B-Table S1-S4**). To validate these results, we used RT-qPCR on a panel of 41 genes spanning over the entire enrichment ratio spectrum, revealing a good correlation with the RNAseq analysis (r^2^=0.405, p<0.0001) (**Figure S1D**). *Calcr* was among the top enriched genes, in line with our previous work implicating CALCR neurons in brain leucine sensing. In addition, IPA pathway analysis revealed activation of p70/S6K and ERK/MAPK signalling pathways, in line with our previous observations implicating these signalling cascades in MBH leucine sensing (**Figure S1E-Table S5**) ^13^.

**Figure 1.**
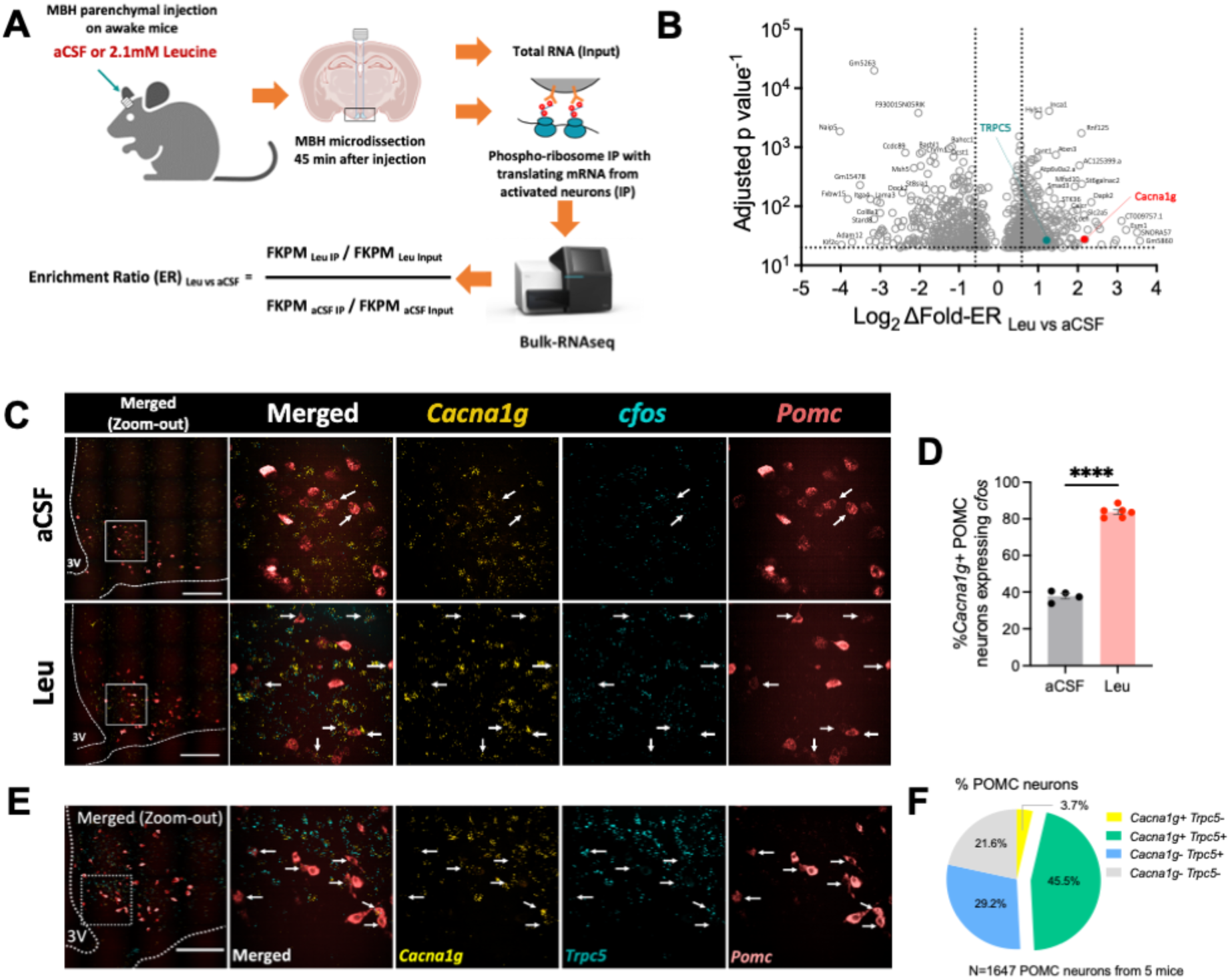
Molecular profiling of hypothalamic leucine-sensing neurons identifies *Cacnag1* as a top marker (**A**) PhosphoTRAP strategy for identification of leucine-responsive neurons in the MBH. (**B**) Volcano plot of differentially enriched markers of MBH leucine-responsive neurons. *Cacna1g* and *Trpc5* are highlighted. (**C-D**) RNAscope fluorescent in situ hybridisation (FISH) of *Cacna1g*, *cfos* and *Pomc* mRNAs in MBH after MBH leucine administration. (**C**) Representative images: white squares: regions shown in the zoom-in images; white arrows: *Cacna1g*+/POMC neurons expressing *cfos;* cale bar: 200 μm; 3V: third ventricle. (**D**) Quantification of imaging analysis. n = 4 for aCSF; n = 6 for Leu. ****p<0.0001. Values are reported as mean ± SEM. (**E-F**) RNAscope FISH of *Cacna1g*, *Trpc5* and *Pomc* mRNAs in MBH. (**E**) Representative images: white square: region shown in the zoom-in images; white arrows: *Cacna1g*+/*Trpc5*+ double positive POMC neurons; scale bar: 200um; 3V: third ventricle. (**F**) Relative proportion of MBH POMC neurons expressing *Cacna1g* and *Trpc5*. n = 1647 POMC neurons from 5 WT mice.

To identify novel markers of leucine-sensing neurons, we focused on transcripts encoding secreted and plasma membrane proteins (**Table S3**) and noticed enrichment in *Cacna1g*, encoding Cav3.1, a T-type calcium channel (**Figures 1B, S1D-S1F**). We chose to further validate this candidate, based on our previous work identifying extracellular calcium entry as an early event required for leucine sensing in POMC neurons ^6^, and work from others indicating that leucine-induced calcium currents contribute to mTORC1 activation ^14^. We used multiplexed RNAscope fluorescence *in situ hybridisation* (FISH) to confirm enrichment of *Cacna1g* in leucine-sensing POMC neurons. As expected, MBH leucine administration significantly increased the proportion of *cfos*^+^ and *cfos^+^/Pomc*^+^ neurons (**Figures S1G-S1H**). Strikingly, 80% of *Cacna1g*^+^/POMC^+^ neurons were activated following MBH leucine administration (**Figures 1C-1D**), confirming that *Cacna1g* expression is enriched in leucine-activated hypothalamic POMC neurons. *Cacna1g* mRNA expression on POMC neurons by RNA-scope was not significantly altered by MBH leucine (**Figure S1I**).

Little is known about the functional relevance of Cav3.1 in POMC neurons, but a recent study suggests that Cav3.1 forms a functional complex with TRPC5 to mediate leptin-induced activation of POMC neurons ^15^. Intriguingly, *Trpc5* is also enriched leucine-activated neurons in our PhosphoTRAP assay (**Figure 1B-Table S3)**. Using RNAscope FISH, we found that ∼50% of POMC neurons express *Cacna1g* and strikingly, ∼90% *Cacna1g*^+^ POMC neurons co-express *Trpc5* throughout the entire rostrocaudal span of the ARH (**Figures 1E-1F**). The colocalization between *Cacna1g^+^*and *Trpc5^+^*was markedly less pronounced in non-POMC neurons (**Figure S1J**). Furthermore, we used HEK293 cells heterologously expressing human Cav3.1 to confirm the direct physical interaction between Cav3.1 and TRPC5, expanding on previous work ^15^ (**Figure S1K**). Thus, Cav3.1 may functionally contribute to the regulation of POMC neuronal activation.

### Cav3.1 activity is required for leucine-induced activation of POMC neurons

To test the hypothesis that Cav3.1 is involved in leucine-induced activation of POMC neurons, we employed TTA-P2, an established pharmacological Cav3.1 inhibitor ^16^. Using our previously established Ca^2+^ imaging setup on acute primary culture of adult hypothalamic neurons ^6^, we found that ∼25% POMC neurons (as marked by EGFP) are activated by acute leucine treatment, but TTA-P2 pre-treatment completely blocked this response (Figures 2A-2D). Likewise, TTA-P2 pre-treatment markedly blunted leucine-induced neuronal activation in human hypothalamic POMC neurons derived from human induced pluripotent stem cells (Figures 2E-2H). Next, we performed brain slice electrophysiological recordings on mouse POMC neurons. In line with previous reports ^5,17^, leucine treatment significantly depolarised POMC neurons, resulting in increased spontaneous action potential firing rate (Figures 2I-2K). Co-treatment with TTA-P2 abolished leucine’s effects (Figures 2L-2N). To test if TRPC5 is also involved in leucine-induced activation of POMC neurons, we used brain slices from *Trpc5* knock-out (KO) mice. *Trpc5* KO abolished leucine-induced depolarisation and increased firing rate of POMC neurons (Figures 2O-2Q), suggesting that Cav3.1 and TRPC5 work as a functional complex to mediate leucine-induced activation of POMC neurons. Last, we tested if Cav3.1 activity is required for the anorectic response to MBH leucine injection in behaving animals using a TTA-P2/Leucine co-injection paradigm (Figure 2R). As expected, MBH leucine suppressed appetite, and this effect was blunted with TTA-P2 co-treatment (Figure 2S). Thus, Cav3.1 activity is required for leucine-induced activation of POMC neurons and appetite suppression.

**Figure 2:**
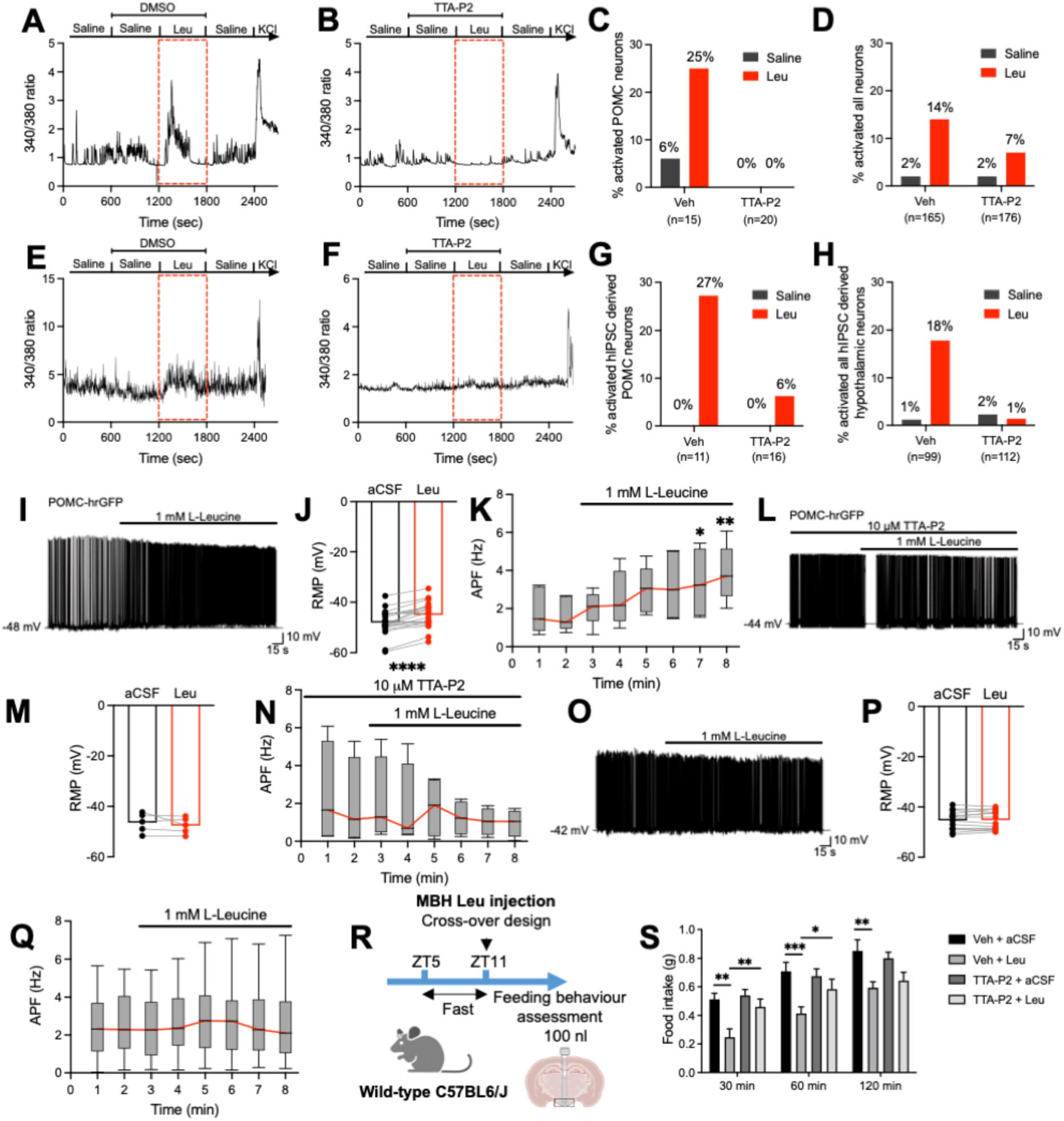
Cav3.1 activity is required for leucine-induced activation of POMC neurons. **(A-D)** Calcium imaging assay on mouse acute primary cultured hypothalamic POMC neurons. Representative calcium trace of a POMC neuron activated by leucine co-treated with vehicle (**A**) and Cav3.1 inhibitor TTA-P2 (**B**). Quantification of activated POMC neurons (**C**) and all cells (**D**) recorded over 4 independent experiments. **(E-H)** Calcium imaging assay on hIPSC derived POMC neurons. Representative calcium trace of a hIPSC derived neuron activated by leucine co-treated with vehicle (**E**) and TTA-P2 (**F**). Quantification of hIPSC derived POMC neuron (**G**) and all cells (**H**) recorded over 4 independent experiments. **(I-K)** Brain slice current-clamp recordings of arcuate POMC neurons with leucine treatment. (**I**) Representative current-clamp trace depicting a characteristic depolarization of arcuate POMC neurons by 1 mM leucine. (**J**) Histogram summarising the acute effect of leucine on the membrane potential of POMC neurons (n=19). (**K**) Box plot of action potential frequency (ARF) in a subpopulation of POMC neurons over time (n=6). **(L-N)** Brain slice current-clamp recordings of arcuate POMC neurons with TTA-P2 pre-treatment prior leucine. (**L**) Representative trace showing that 1 mM leucine fails to induce a depolarization of POMC neurons exposed to 10 μM TTA-P2. (**M**) Histogram summarizing the acute effect of leucine on the membrane potential of POMC neurons in the presence of 10 μM TTA-P2 (n=5). (**N**) Box plot of APF of POMC neurons in the presence of TTA-P2 over time (n=4). **(O-Q)** Brain slice current-clamp recordings of arcuate POMC neurons from *Trpc5* KO mice with leucine treatment. (**O**) Representative trace showing that 1 mM leucine fails to induce a depolarization of POMC neurons on a *Trpc5* KO background. (**P**) Histogram summarizing the acute effect of leucine on the membrane potential of POMC neurons deficient for TRPC5 (n=14). (**Q**) Box plot of APF of POMC neurons deficient for TRPC5 over time (n=13). **(R-S)** Feeding responses assessment after MBH leucine injection on awake mice. (**R**) Diagram of the experimental paradigm. (**S**) Acute feeding response after MBH leucine co-treated with vehicle or TTA-P2. n = 10 per group. *p<0.05, **p<0.01, ***p<0.001, ****p<0.0001. Range bars in box plots (K, N, Q) indicates min to max and comparisons were made versus the minute bin prior leucine treatment. Values in (S) are reported as mean ± SEM.

### Cav3.1 is a leucine sensor

To test the possibility that leucine might directly interact with Cav3.1 and modulate its function, we used HEK293 cells stably overexpressing human Cav3.1 (HEK293-hCav3.1). In HEK293-hCav3.1 cells loaded with cell-permeable Ca^2+^ sensitive Fluo8-AM dye, we confirmed dose-dependent activation of hCav3.1 by potassium chloride (KCl)-induced depolarisation (**Figures S2A-S2D**). Leucine treatment significantly activated hCav3.1 and enhanced KCl-induced hCav3.1 activation (Figures 3A-3B). These effects were not observed in wild-type HEK293 cells (**Figures S2E-S2F**), suggesting that leucine has a direct effect on Cav3.1 function. Activation of hCav3.1 is specific to leucine among all branched-chain amino acids (Figures 3C-3D), and is mTOR independent (Figures 3E-3F). To elucidate the electrophysiological mechanisms through which leucine affects Cav3.1 function, we performed whole-cell voltage-clamp recordings on HEK293-hCav3.1. In control cells, the activation (V50_act_Con_: −41.78 ± 12.83 mV) and inactivation curves (V50_act_Con_: −47.77 ± 11.91 mV) were comparable with the published literature ^18^. Leucine treatment produced a significant left-shift of both the activation (V50_act_Leu_: −57.42 ± 11.18 mV) and inactivation curves (V50_inact_Leu_: −60.50 ± 11.01 mV) (Figures 3G-3J). The resting membrane potential of wild-type HEK293 cells has been reported to be at about −30 to −50 mV ^19^. Importantly, according to our data, half of hCav3.1 are activated at this voltage band in vehicle-treated cells, while with leucine treatment hCav3.1 is fully activated (Figures 3I).

**Figure 3:**
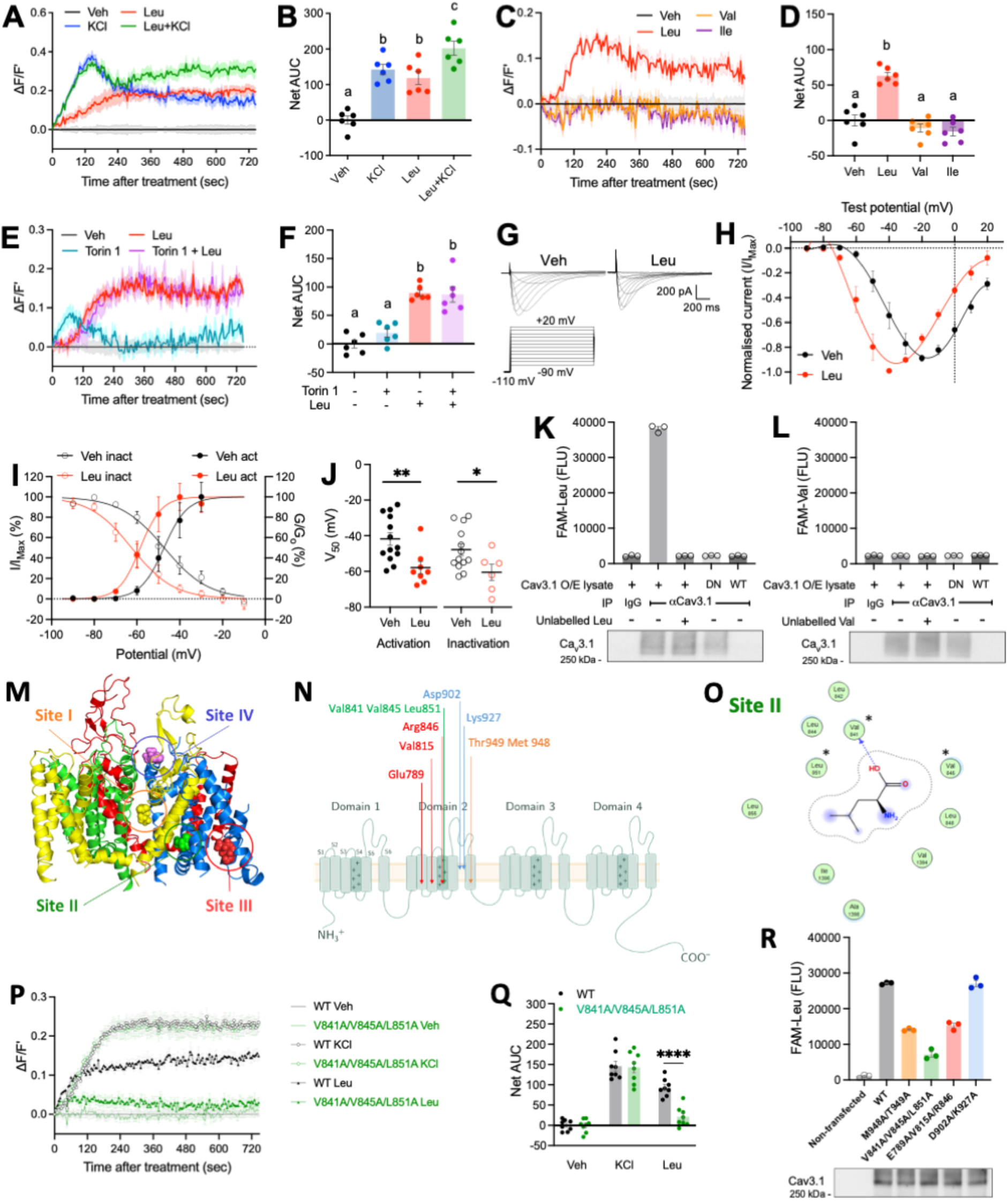
Cav3.1 is a leucine sensor. **(A-F)** Fluo8 calcium flux assay of HEK293-hCav3.1 cells. (**A-B**) Calcium responses and AUC quantification after KCl and Leu treatments. (**C-D**) Calcium responses and AUC quantification after Leu, Val and Ile treatments. (**E-F**) Calcium responses and AUC quantification after Leu treatment with or without mTOR inhibitor Torin 1 (100 nM) pre-treatment. n = 6 per group merged from 2 independent experiments with triplicates each. (**G-J)** Whole-cell patch-clamp recording of HEK293 transiently expressing hCav3.1 treated with leucine. (**G**) Representative traces of Ca^2+^ currents recorded in vehicle solution (left) and after application of 1 mM leucine. In both cases, cells were held at −100 mv and increasing steps of 10 mV were applied from −90 to +20 mv. (**H**) Normalised current-voltage (I-V) curves of HEK293-hCav3.1 cells treated with vehicle or leucine under the activation protocol. Stead-state activation (G/G_max_) and inactivation (I/I_max_) curves (**I**) and V_50_ histogram (**J**) of HEK293-hCav3.1 cells treated with vehicle or leucine. Veh_act: n=13, Leu_act: n=8, Veh_inact: n=13, Leu_inact: n=6. **(K-L)** In vitro binding assay of hCav3.1 to fluorescent labelled leucine (**K**) and valine (**L**). Post-assay Western blots below the plots validate successful immunoprecipitation (IP). IgG: unimmunized rabbit IgG control; DN: heat-denatured cell lysate prior IP; WT: cell lysate from non-transfected HEK293 cells. Data represent 3 technical replicates upon fluorescence measurements. These experiments has been repeated twice with similar results. Data of representative sets of experiment are shown. **(M)** Human Cav3.1 protein (PDB id: 6KZO) is shown in cartoon representation with domain I, II, III and IV coloured in red, light blue, yellow and green, respectively. The representative binding poses of L-leucine are shown in sphere representation with poses relevant to different sites coloured distinctly: yellow (putative site I, marked with an orange circle), green (putative site II, marked with a green circle), red (putative site III, marked with a red circle) and light pink (putative site IV, marked with a blue circle). The figure was generated in PyMol 2.5 (Schrodinger LLC). **(N)** Schematic representation of hCav3.1 critical amino acid residues of corresponding predicted leucine binding sites chosen for mutagenesis experiments, colour-coded as in (**M**). **(O)** 2D ligand interactions diagrams for the docked poses of leucine at Site II. The residues mutated for experimental validation are shown with asterisks. The dotted arrow signs indicate hydrogen bonding whilst the dotted contour around the ligand pose indicate hydrophobic interactions with residues shown. These diagrams were generated in MOE. **(P-Q)** Fluo8 calcium flux assay of HEK293-hCav3.1 Site II mutant (V841A/V845A/L851A) cells. Calcium responses (**P**) and AUC quantification (**Q**) after KCl and Leu treatments. n=8 merged from 4 independent experiments with duplicates each. **(R)** In vitro fluorescent leucine binding assay of WT and Site I–IV hCav3.1 mutants. Post-assay Western blot below the plot validates successful IP. Data represent 3 technical replicates upon fluorescence measurements. These experiments have been repeated twice with similar results. Data of a representative experiment is shown. Groups denoted with different letters in (B, D, F) indicate significant difference (p<0.05). *p<0.05, **p<0.01, ****p<0.0001. Values and calcium response traces in (A, C, E) are reported as mean ± SEM.

Given that mTOR signalling is unlikely to mediate leucine-induced Cav3.1 activation (Figures 3E-3F) ^6^, and that Cav3.1 phosphorylation does not change its voltage dependence ^18^, we reasoned that leucine might directly bind to Cav3.1 to modulate its activity. To test this idea, we adopted an immunoprecipitation-based *in vitro* leucine binding assay, previously used to test the leucine-binding property of the leucine sensor Sestrin2 ^20^. Using fluorescently-labelled leucine (FAM-leucine), we detected marked binding of FAM-leucine with hCav3.1, out-competed following incubation with excess amounts of unlabelled leucine, indicating that leucine can directly bind to hCav3.1 (Figure 3K). Moreover, heat-denaturation of hCav3.1 abolished FAM-leucine binding, indicating that intact protein folding of hCav3.1 is needed for leucine binding (Figure 3K). Of note, FAM-valine failed to bind to hCav3.1, suggesting a leucine-specific binding mechanism (Figure 3L).

To identify leucine’s binding site on hCav3.1, we performed a domain mapping experiment by generating a series of non-overlapping truncated hCav3.1 based on its 4 distinct transmembrane domains, and tested them using the FAM-leucine binding assay. Unexpectedly, none of the 4 domains alone bound leucine to a comparable extent of the full-length hCav3.1 (**Figures S2G-S2I**), implying that the leucine binding site may involve intact 3D structure. We next performed an *in silico* leucine docking analysis using the recently cryo-EM-resolved 3D structure of the human Cav3.1 protein (PDB id: 6KZO)^21^. We first performed blind docking of L-leucine as the active ligand and with L-valine as a decoy against the entire hCav3.1 structure. After further focused docking, we shortlisted four plausible leucine binding sites for experimental validation (hereafter termed Site I-IV) (Figures 3M-3O, S2M, S2P and S2S). To test the importance of these sites in leucine binding and effects on leucine-induced hCav3.1 activation, we performed site-directed mutagenesis of the critical residues on these sites to alanine (Figure 3N), generated HEK293 stable cell lines expressing these mutated hCav3.1 channels and used Western blot to confirm their expression (**Figures S2L**), and then investigated these using the Fluo8 Ca^2+^ flux and *in vitro* FAM-leucine binding assays. The Site IV mutant (D902A/K927A) did not affect the channel function (**Figures S2T-S2U**) nor its binding to FAM-leucine (Figure 3R). Site I (M948A/T949A) and Site III (E789A/V815A/R846A) mutants did not respond to KCl or leucine (**Figures S2N-S2O-S2Q-S2R**), and had markedly impaired leucine binding (Figure 3R), suggesting major alterations to the protein structure affecting both normal voltage-sensing and leucine binding. In contrast, the Site II mutant (V841A/V845A/L851A) showed comparable response to KCl as WT, but leucine-induced Ca^2+^ influx was significantly blunted (Figures 3P-3Q), and binding affinity to FAM-leucine was the lowest (∼-75% vs. WT) (Figure 3R). Thus, Site II-a cytoplasm-facing hydrophobic pocket formed by the two voltage-sensing S4 segments of domain II and III-is a functionally critical leucine binding site. Additionally, amino acid sequence alignments show that these two S4 segments of hCav3.1 are fully conserved among mammals but not with zebrafishes nor with hCav3.2 and hCav3.3 (**Figure S2V**), suggesting Cav3.1-mediated leucine sensing mechanism may be evolutionally conserved in mammals.

Previously, it has been demonstrated that mTOR signalling is required for the appetite-suppressive effects of MBH leucine *in vivo* ^8^, and work from others showed that Ca2+ influx activates mTOR in a calmodulin/hVps34 dependent manner ^14^, which are ubiquitously expressed and has been detected in human hypothalamus and HEK293 cells (Human Protein Atlas). Accordingly, to test the prediction that leucine can activate mTOR signalling via Cav3.1, we used HEK293-Cav3.1 cells, which show robust Thr389 phosphorylation or S6K in response to leucine treatment. TTA-P2 significantly blunted leucine-induced activation of the mTOR signalling pathway (**Figures S2J-S2K**), suggesting that Cav3.1 contributes to leucine-induced mTOR activation. Collectively, these data support a role for Cav3.1 as a leucine sensor mediating leucine-induced activation of POMC neurons and mTOR signalling.

### Mediobasal hypothalamic *Cacna1g* is required for the appetite and weight-suppressing effect of dietary proteins

To investigate the contribution of hypothalamic Cav3.1 in the regulation of energy homeostasis we developed an AAV-based CRISPR mediated knock-out of *Cacna1g* in the MBH (hereafter termed Cacna1g^MBH^ ^KO^) (Figures 4A**, S3A-S3C**), and validated expected indel mutations and top predicted off-target genes in mouse hypothalamic mHypoE-N46 cells and hypothalamic tissues (Figures 4D**, S3D-S3E**). Western Blot analysis using an anti-Cav3.1 C-terminal specific antibody revealed a 64% KO efficiency in the MBH of Cacna1g^MBH^ ^KO^ mice (Figures 4B-4C).

**Figure 4:**
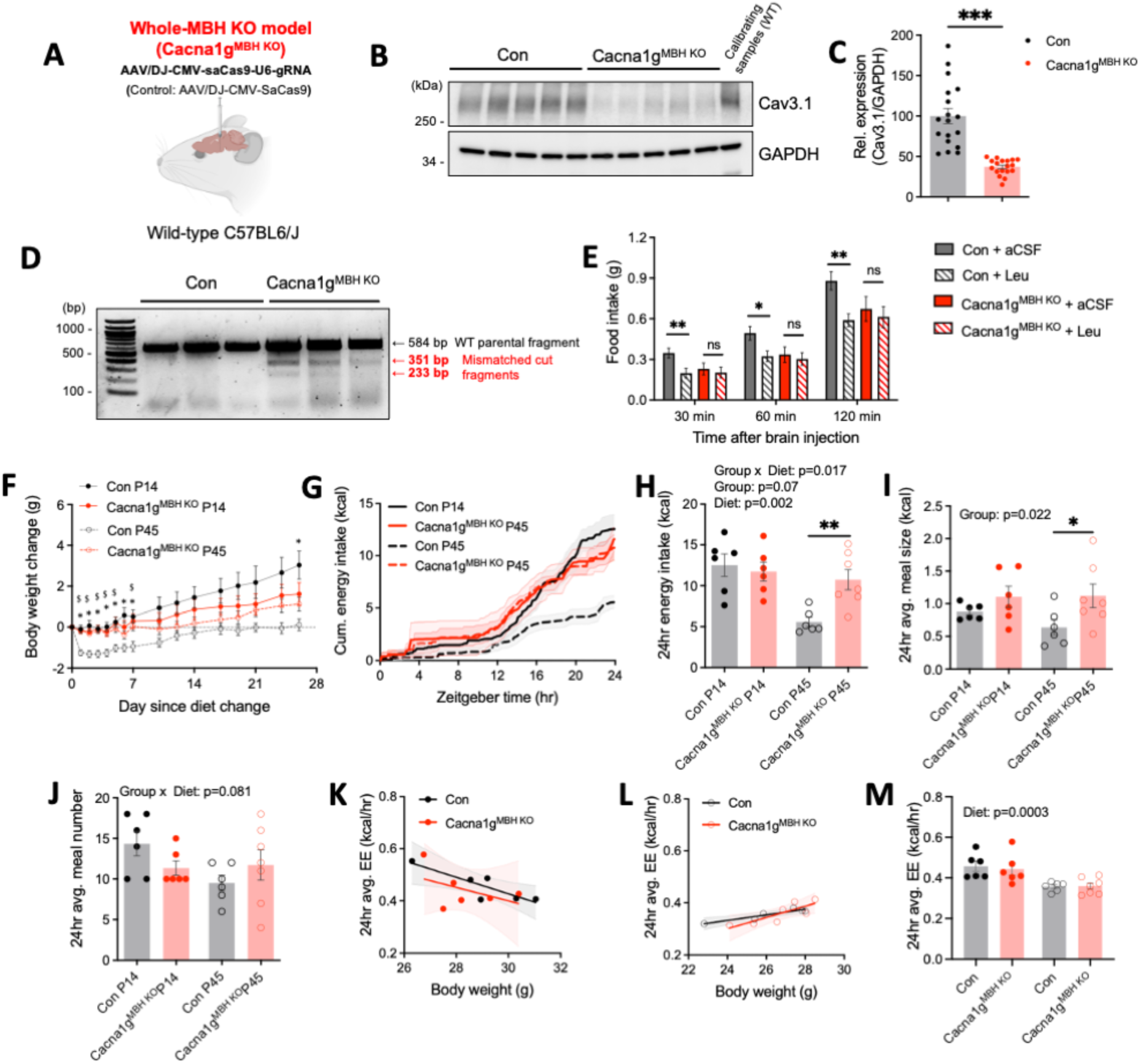
Mediobasal hypothalamic *Cacna1g* is required for the appetite- and weight-suppressing effect of dietary proteins. **(A)** Diagram of experimental strategy to generate Cacna1g^MBH^ ^KO^ mice. **(B-C)** Western blot analysis of the CRISPR knockout efficiency of Cav3.1 in Cacna1g^MBH^ ^KO^ mediobasal hypothalamus with a C-terminal specific Cav3.1 antibody. GAPDH from the same blot was used as a loading control. Representative image (**B**) and densitometric quantification (**C**) of the blots. n = 18 per group. **(D)** Representative DNA electrophoresis image of T7 endonuclease I mutation detection assay of genomic DNA isolated from the MBH tissues of Con and Cacna1g^MBH^ ^KO^ mice. Black arrow indicates parental unmodified PCR amplicons and red arrows indicate digested mutated fragments of expected size. **(E)** Acute feeding response after MBH leucine injection in Con and Cacna1g^MBH^ ^KO^. Con: n = 11; MBH KO: n=12. **(F)** Body weight change of Con and Cacna1g^MBH^ ^KO^ over the course of 4 weeks feeding on 14% normal protein (P14) diet and 45% hight protein (P45) diet. **(G-J)** Feeding behaviour assessments of Con and Cacna1g^MBH^ ^KO^ mice on P14 and P45 diets during indirect calorimetry measurement. (**G**) 24-hr profile of cumulative energy intake, (**H**) 24-hr total energy intake, (**I**) 24-hr average meal size and (**J**) 24-hr average meal number. **(K-M)** Energy expenditure of Con and Cacna1g^MBH^ ^KO^ mice on P14 and P45 diets during indirect calorimetry measurement. ANCOVA analysis of average 24-hr energy expenditure against body weight of Con and Cacna1g^MBH^ ^KO^ on P14 (**K**) and P45 (**L**) diets, 24-hr average energy expenditure (**M**). For (**F-M**) Con-P14: n = 6; Con-P45: n = 6; KO-P14: n = 7; KO-P45: n = 7. *p<0.05, **p,0.01, ***p<0.001. Values are reported as mean ± SEM.

Consistent with the effect of MBH pharmacological inhibition of Cav3.1 with TTA-P2, MBH leucine failed to reduce food intake in Cacna1g^MBH^ ^KO^ mice (Figure 4E), indicating that Cav3.1 activation is required for leucine-induced anorexia. To determine if leucine sensing through Cav3.1 contributes to the feeding and metabolic responses to high-protein diets, we transitioned Cacna1g^MBH^ ^KO^ and control mice to either a 45% protein diet (high protein content, hereafter termed P45) or a 14% protein diet (normal protein content, hereafter termed P14) (**Table S6**). Over the course of 4 weeks, P45 feeding significantly reduced weight gain in control mice, but this effect was blunted in Cacna1g^MBH^ ^KO^ mice (Figure 4F). Decreased weight gain in the control group under P45 feeding was accounted for by a significant reduction in energy intake during the active phase through a specific decrease in meal size, which failed to occur in Cacna1g^MBH^ ^KO^ mice (Figures 4G-4J). P45 feeding significantly reduced energy expenditure but no difference was observed between Cacna1g^MBH^ ^KO^ and control groups (Figures 4K-4M, S4A-S4F). We did not detect overt differences in respiratory exchange ratio (RER) and locomotor activity among the control and Cacna1g^MBH^ ^KO^ mice under both feeding paradigms (**Figures S4B, S4C, S4G-S4H)**. Together, these data revealed MBH Cav3.1 mediates protein-induced satiation and is required for the weight-suppressive effect of high protein feeding.

To further investigate the role of MBH Cav3.1 in metabolic heath, we assessed glucose homeostasis in Cacna1g^MBH^ ^KO^ mice. We detected a non-significant trend towards improved glucose tolerance in Cacna1g^MBH^ ^KO^ mice under P14 feeding (**Figure S4D**), which was not observed during P45 feeding (**Figure S4I**). Insulin sensitivity was comparable between groups under both P14 and P45 feeding **(Figures S4E-S4J**). Post-absorptive plasma amino acid concentrations and urinary markers of nitrogen disposal were markedly affected by high protein feeding but did not overtly differ between groups (**Figures S4K-S4R and Table S7**). Overall, MBH Cav3.1 does not appear to impinge on glucose metabolism regulation and peripheral amino acid availability control.

Previous report showed that whole-body *Cacna1g* KO protects against diet-induced obesity (DIO) ^22^, but the underpinning mechanisms are unclear. To elucidate the role of MBH Cav3.1 in DIO, we maintained Cacna1g^MBH^ ^KO^ mice on 60% high-fat diet and assessed their metabolic phenotypes. Under these conditions, weight gain (**Figures S5A-S5C**), energy intake (**Figures S5D-S5G**) and expenditure (**Figures S5H-S5J**), locomotor activity **(Figures S5K**) and RER (**Figures S5L**) did not overtly differ between Cacna1g^MBH^ ^KO^ and WT controls. However, post-mortem RT-qPCR analysis revealed that MBH *Cacna1g* expression was significantly downregulated following 8-weeks HFD feeding in control mice **(Figures S5M**), which might explain the lack of phenotypic difference between Cacna1g^MBH^ ^KO^ and WT mice under these conditions. Overall, these data imply that MBH Cav3.1 is not involved in the protective effects of whole-body Cacna1g KO against DIO, and suggest that HFD feeding may blunt Cav3.1-mediated leucine sensing in the MBH.

### *Cacna1g* in hypothalamic POMC neurons is required for the appetite-suppressing effect of dietary proteins

To test if the feeding phenotype observed in Cacna1g^MBH^ ^KO^ mice is mediated via Cav3.1 activity in ARH POMC neurons, we used a CRISPR strategy to knockout *Cacna1g* specifically in ARH POMC neurons by stereotactical injection of gRNAs-expressing AAV to the MBH of transgenic mice with SpCas9 expression exclusively in POMC neurons (hereafter termed Cacna1g^POMC^ ^KO^, Figures 5A**, S3B-S3C**). To establish an immunofluorescence-based KO validation strategy, we first validated an anti-Cav3.1 C-terminal specific antibody using the global Cacna1g KO mice ^23^. As expected, we detected robust Cav3.1 expression in the MBH and thalamus (**Figures S3F-G**). We then used immunofluorescence with this antibody to validate POMC-specific KO of *Cacna1g* and observed a ∼60% reduction in Cav3.1 expression in mCherry-expressing transduced POMC neurons in Cacna1g ^POMC^ ^KO^ mice (Figures 5B-5C-S6A-S6B).

**Figure 5:**
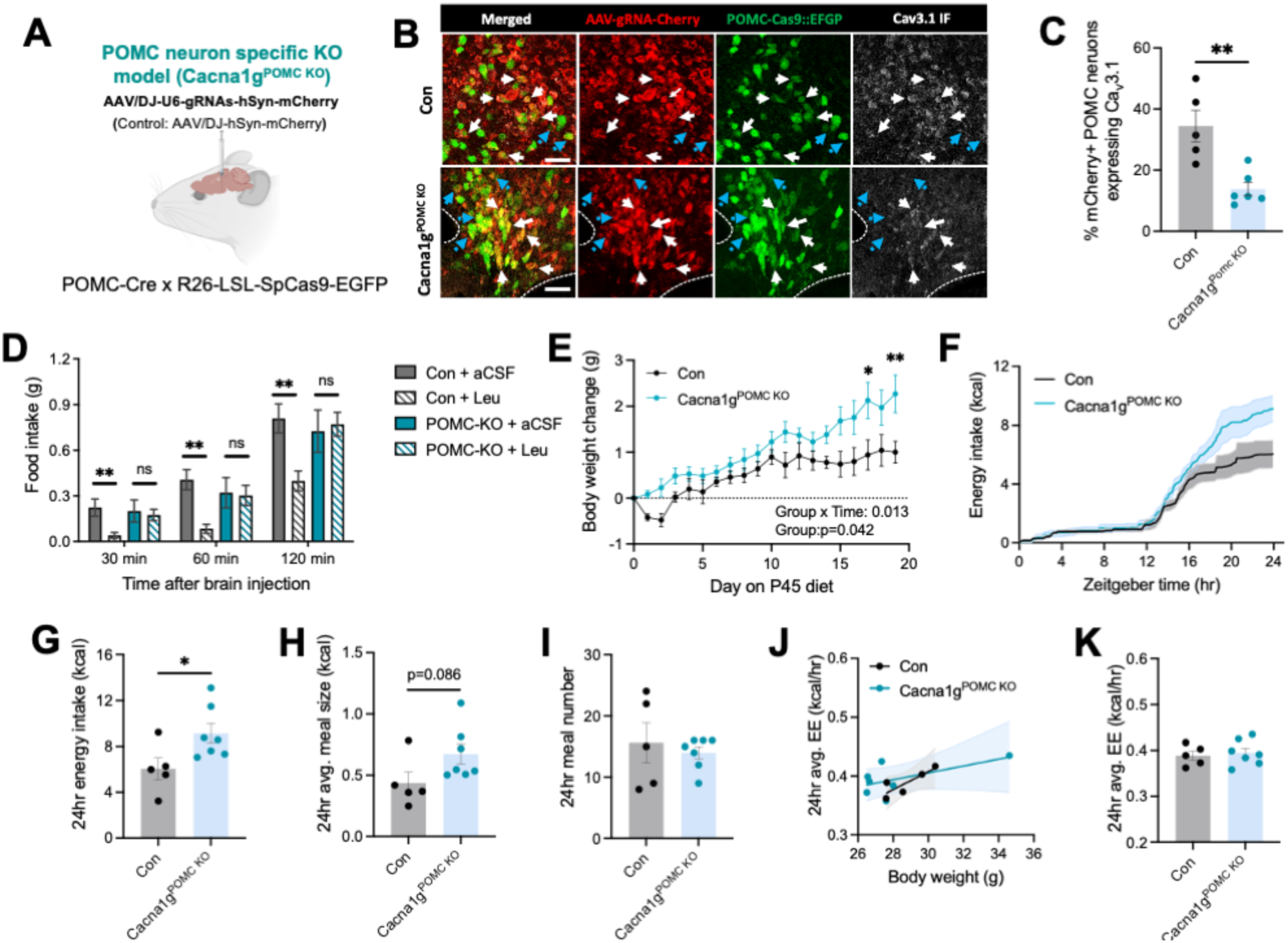
*Cacna1g* in hypothalamic POMC neurons is required for the appetite-suppressing effects of high protein diet feeding. **(A)** Diagram of experimental strategy to generate Cacna1g^POMC^ ^KO^ mice. **(B-C)** Confocal **i**mmunofluorescence (IF) microscopy analysis of the CRISPR knockout efficiency of Cav3.1 in Cacna1g^POMC^ ^KO^ mediobasal hypothalamus with a C-terminal specific Cav3.1 antibody validated in Fig. S3F. (**B**) Zoom-in images of the regions highlighted with white squares in (Fig. S6A). White arrows with solid line: example POMC+/mCherry+/Cav3.1+ neurons. Blue arrows with broken line: example POMC+/mCherry+/Cav3.1-neurons. Scale bar: 50 µm. (**C**) Quantification for % mCherry+ transduced POMC neurons expressing Cav3.1. Con: n = 5; Cacna1g^POMC^ ^KO^: n = 6. **(D)** Acute feeding response after MBH leucine injection in Con and Cacna1g^POMC^ ^KO^. Con: n = 10; KO: n=8. **(E)** Body weight change of Con and Cacna1g^POMC^ ^KO^ over the course 3 weeks feeding on P45 diet. **(F-I)** Feeding behaviour assessments of Con and Cacna1g^POMC^ ^KO^ mice on P45 diets during indirect calorimetry measurement. (**F**) 24-hr profile of cumulative energy intake, (**G**) 24-hr total energy intake, (**H**) 24-hr average meal size and (**I**) 24-hr average meal number. **(J-K)** Energy expenditure of Con and Cacna1g^POMC^ ^KO^ mice on P45 diets during indirect calorimetry measurement. (**J**) ANCOVA analysis of average 24hr energy expenditure against body weight of Con and Cacna1g^MBH^ ^KO^ on P45 diets, (**K**) 24hr average energy expenditure. For (E-K), Con: n = 5; Cacna1g^POMC^ ^KO^: n = 7. *p<0.05, **p,0.01. Values are reported as mean ± SEM

POMC-specific KO of *Cacna1g* was sufficient to blunt the anorectic effect of MBH leucine injection (Figure 5D). During maintenance on P45, Cacna1g^POMC^ ^KO^ mice gained significantly more weight compared to controls (Figure 5E), which was mediated by an increase in energy intake during the active dark phase (Figures 5F-5G) and a trend towards increased meal size but no difference in meal number (Figures 5H-5I). Energy expenditure (Figures 5J**-S6C**), locomotor activity (**Figure S6D**), RER (**Figure S6E**), glucose tolerance (**Figure S6F**) and urinary parameters (**Figures S6G-S6I**) did not differ between groups. Thus, Cacna1g^POMC^ ^KO^ mice broadly phenocopied observations seen in Cacna1g^MBH^ ^KO^ mice during high protein diet feeding, supporting the idea that it is Cav3.1 expressed in POMC neurons mediating high protein diet feeding responses.

### Pharmacological activation of hypothalamic Ca_v_3.1 reduces appetite and promotes weight loss in lean mice

We explored the potential of Cav3.1 as a weight loss target using SAK3, a novel small molecule pharmacological activator of Cav3.1 ^24^. MBH injection of SAK3 reduced 24-hr food intake and body weight gain in a dose-dependent manner (Figures 6A-6B-S6J). This effect is lost in Cacna1g^MBH^ ^KO^ mice (Figures 6C-6F), confirming that Cav3.1 activation mediates SAK3 action.

**Figure 6:**
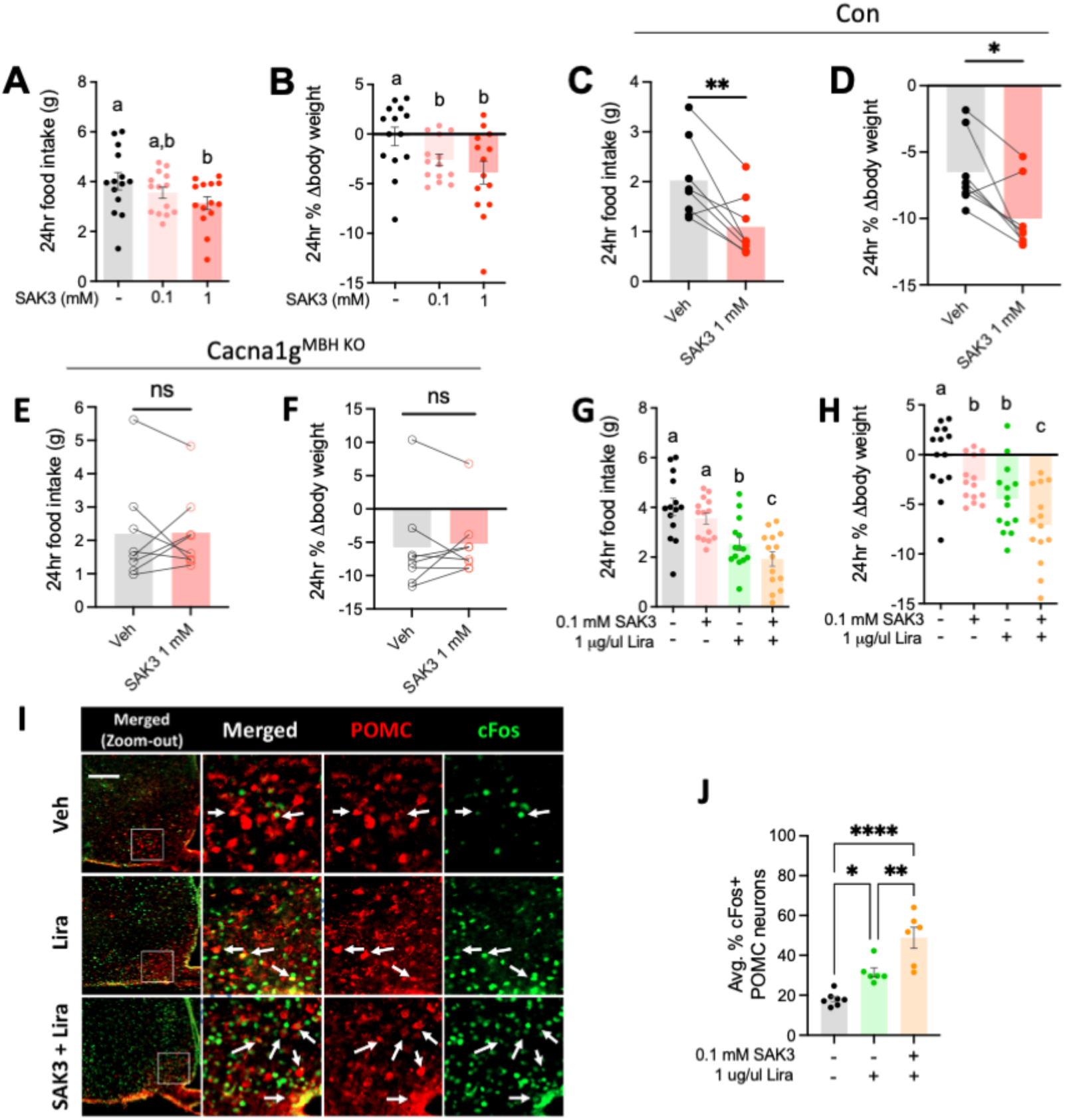
Pharmacological activation of hypothalamic Cav3.1 reduces appetite and promotes weight loss in lean mice. **(A-B)** 24-hr feeding responses (**A**) and % body weight changes (**B**) after MBH injection of Veh, 0.1 mM and 1 mM Cav3.1 activator SAK3. n = 14 per group. **(C-F)** Hypothalamic expression of Cav3.1 is required for the feeding effects of SAK3. 24-hr feeding responses and % body weight changes after MBH injection of Veh and 1mM SAK3 in Con (n = 6) (**C-D**) and Cacna1g^MBH^ ^KO^ mice (n = 8) (**E-F**). **(G-H)** 24hr feeding responses (**G**) and % body weight changes (**H**) after MBH injection of Veh, 0.1 mM SAK3, 1 μg/μl liraglutide (Lira) and SAK3+Lira co-treatment. n = 14 per group. **(I-J)** Immunofluorescence (IF) microscopy analysis of POMC neuron activation after Lira and SAK3+Lira. (**I**) Representative fluorescence images of POMC (red) and cFos (green). White squares: regions shown in the zoom-in images. White arrows: neurons double positive for POMC and cFos IF. Scale bar: 200um. (**J**) Quantification of images analysis. Groups denoted with different letter in (A, B, G, H) indicate significant difference (p<0.05). *p<0.05, **p<0.01, ****p<0.0001. Values are reported as mean ± SEM.

Hypothalamic POMC neurons has been shown to mediate the anorectic action of the FDA-approved weight loss drug liraglutide, a GLP-1 receptor agonist where TRPC5 expression is required ^25,26^. Thus, we next tested the effect of a subthreshold dose of SAK3 on liraglutide-induced appetite- and weight-suppression. As predicted, MBH SAK3 increased the anorectic and weight-suppressive effect of liraglutide (Figures 6G-6H-S6K). Immunohistochemistry imaging analysis confirmed that SAK3 increased liraglutide-induced activation of POMC neurons under these conditions (Figures 6I-6J). Likewise, subthreshold SAK3 increased the appetite- and weight-suppressive action of leptin and the 5HT-receptor 2C agonist lorcaserin, which both signal through POMC neurons to regulated appetite and energy balance (**Figures S6L-S6Q**). Thus, MBH Cav3.1 plays a role in integrating satiety signals *in vivo* and might represent an efficacious weight loss target.

### Intranasal administration of Cav3.1 activator as an anti-obesity treatment

We tested the potential weight loss benefits of MBH Cav3.1 activation with SAK3 in diet-induced obese mice using a non-invasive drug delivery method to the central nervous system via the intranasal (IN) route (Figure 7A) ^26^. In a pilot experiment, we found that IN delivery of SAK3 induced weight loss in lean mice maintained on chow diet overnight (Figure 7B), with no obvious adverse effects. We developed a mass spectrometry-based method to detect SAK3 abundance in tissues and confirmed presence of SAK3 in the MBH 1 hour after IN SAK3 dosing (Figure 7C), demonstrating the feasibility of using the IN route to deliver SAK3 to the MBH. We could also detect SAK3 in heart tissues but at concentrations ∼13-fold lower than in the MBH (within the femtomolar range) (Figure 7D). Next, we tested the weight loss effect of IN SAK3 in diet-induced obese mice (Figure 7E). After 12-weeks of HFD feeding, all mice were obese and weighed over 40g. Daily IN SAK3 dosing for 2 weeks significant reduced food intake (Figure 7F), leading to significant weight loss, and significantly increased the appetite- and weight-suppressive action of peripherally-administered liraglutide (Figures 7F-7H). Collectively, these results support the feasibility of targeting MBH Cav3.1 via the intranasal route as a weight loss strategy.

**Figure 7:**
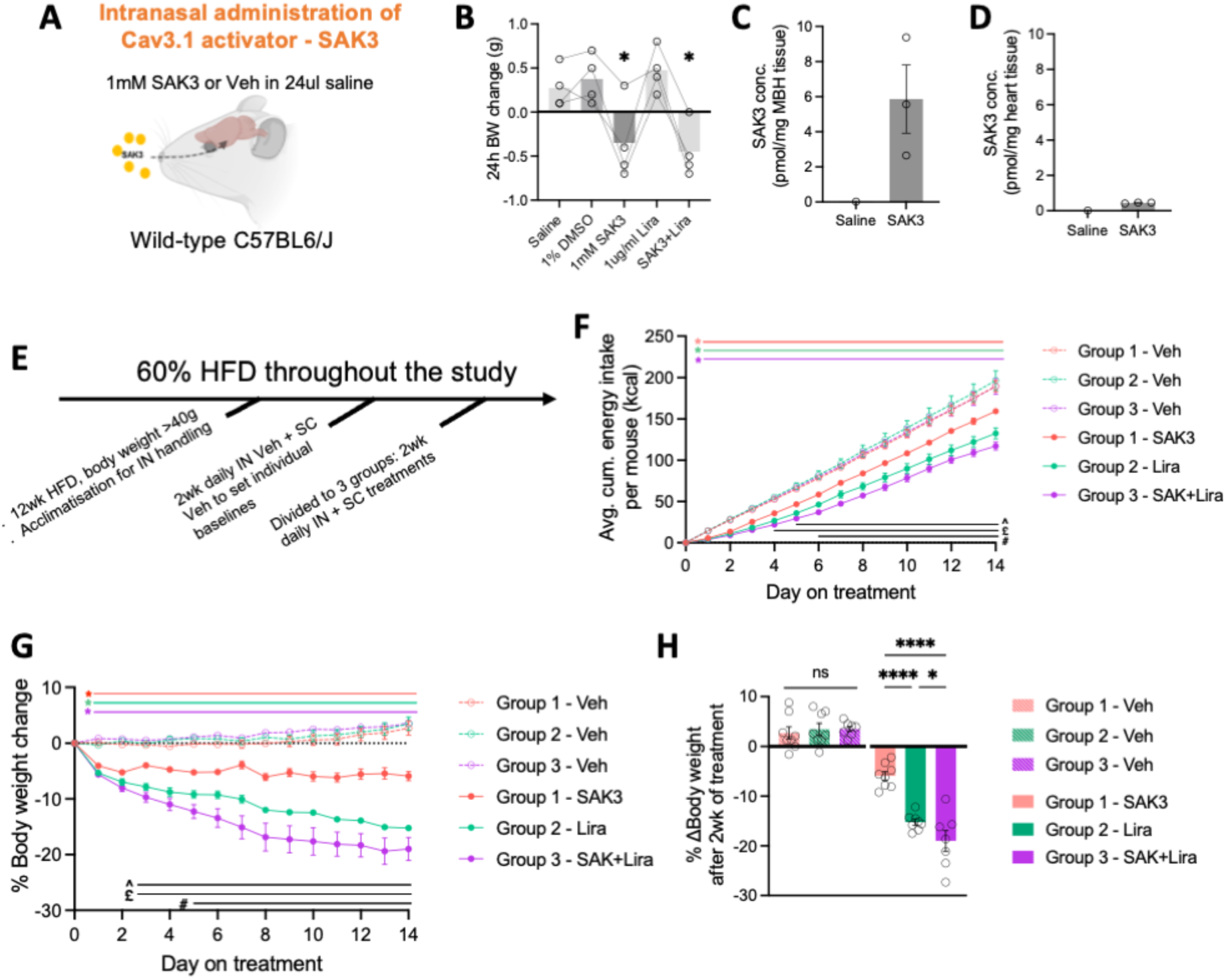
Intranasal administration of Cav3.1 activator as an anti-obesity treatment. **(A)** Diagram of experimental strategy using intranasal administration (IN) of SAK3 to activate hypothalamic Cav3.1 activity. **(B)** 24 hours body weight change after IN administration of indicated substances on lean mice fed on normal chow. Comparisons were made against saline treated condition. Each treatment were 96 hours apart. n = 4 per repeated testing condition. **(C-D)** Mass spectrometry quantification of the tissue abundance of SAK3 in the MBH (**C**) and heart (**D**) 1 hour after IN administration. n = 1 for IN saline; n = 3 for IN SAK3. **(E)** Diagram of the experimental paradigm to test the weight loss effect of IN SAK3 in high fat diet induced obesity mouse model. n = 8 per group for (H-F). **(F)** Cumulative energy intake since the start of respective treatments over the course of 2 week treatment. **(G-H)** % Body weight change normalised to the start of respective treatments over the course of 2-week treatment (**G**). Histogram summarising the terminal % body weight change (**H**). For (F-G), on the top over the plots, colour-coded * indicates p<0.05 between respective Drug vs Veh treatment phases within the same group; at the bottom of the plots, ^p<0.05 SAK3 vs Lira; ^£^p<0.05 SAK vs SAK3+Lira; ^#^p<0.05 Lira vs SAK3+Lira. *p<0.05, ****p<0.0001. Values are reported as mean ± SEM.

## Discussion

In this study, we uncover an unexpected role for Cav3.1 as a neuronal leucine sensor mediating the anorectic action of hypothalamic leucine sensing and high protein feeding. We found that *Cacna1g* expression is highly enriched in MBH neurons activated by leucine, about 4 folds over non-responsive neurons. Subsequent experiments revealed that *Cacna1g* is not only a marker for leucine-sensing neurons, but also functionally required for the rapid expression of calcium currents in response to increased leucine levels. *Ex vivo* patch-clamp recordings confirmed that Cav3.1 is required for leucine-induced activation of MBH POMC neurons. We used hCav3.1-expressing HEK293 cells to distinguish the role of Cav3.1 as a generic gating mechanism or a specific mediator of leucine-induced neuronal activation. Surprisingly, acute leucine exposure activates Cav3.1 via significantly enhancing its voltage sensitivity and thus lowering the activation threshold of the channel. In line with our previous report ^6^, we found that mTOR signalling is not required for the acute leucine-induced activation of Cav3.1. This is consistent with the observation that while current density of Cav3.1 can be regulated through phosphorylation on multiple phosphorylation sites, none of Cav3.1 phosphorylation sites are able to shift its voltage-dependence, as observed after leucine treatment ^18^. Instead, we find that leucine sensing through Cav3.1 contributes to leucine-induced mTOR activation, identifying a previously unappreciated mechanism of regulation for this master growth controller.

The acute nature of leucine-induced activation of Cav3.1 prompted us to reason that a more direct mechanism may be involved. Recently, GABA (γ-Aminobutyric acid)-a modified amino acid neurotransmitter – has been shown to directly bind to voltage-gated potassium channels KCQN3 and 5 ^28^. By analogy, we hypothesised that leucine-induced Cav3.1 activation might also occur via direct binding of leucine to the channel. Combining *in silico* analyses, *in vitro* binding, and functional assays, we revealed that leucine (but not valine or isoleucine) binds to the hydrophobic pocket located between the voltage-sensing S4 segments of domain II and III of Cav3.1, linking leucine’s action to Cav3.1 voltage sensing. Importantly, the leucine sensing site on Cav3.1 is fully conserved among mammals but not among other vertebrate groups. Among the three members of T-type VGCCs, there is a marked variation in the sequence of the amino acid residues involved in leucine binding, suggesting that the leucine-sensing property of Cav3.1 might be unique among VGCCs, although further study will be needed to confirm this prediction. Altogether, these results suggest that Cav3.1 is a neuronal leucine sensor shared among mammalian species.

The mechanism through which leucine activates Cav3.1 is specific among the three BCAAs. This is in line with our previous study that MBH valine injection does not alter feeding behaviour nor activate hypothalamic POMC neurons ^5,6^. To date, the only described cell-intrinsic leucine-specific sensing mechanism is Sestrin1/2-mediated mTORC1 activation ^20,29^. Unlike Cav3.1, Sestrins-mTORC1 signalling is ubiquitously expressed (Human Protein Atlas). Acute hypothalamic activation of mTORC1 signalling results in anorexia and has been shown to contribute to MBH leucine’s anorectic effects ^4,8^. A recent study has demonstrated that ARH POMC neurons respond to mTORC1 signalling heterogeneously, depending on their glutamatergic or GABAergic molecular identity, but collectively orchestrate anorexia ^30^. *Cacna1g* expressing POMC neurons are heterogenous, ∼40% are glutamatergic (*Slc17a6* expressing; encoded for VGLUT2) and ∼60% are GABAergic (*Gad1* expressing) ^31^, thus it is expected that their neuronal responses to mTORC1 activation might be diverse. Interestingly, we demonstrate that Cav3.1-induced calcium currents activate mTORC1 signalling, presumably via a previously described calmodulin-hVps34 mechanism ^14^. Taken together, these observations suggest a multiphasic sequence of leucine-induced activation of anorectic POMC neurons: leucine acutely activates Cav3.1 which causes the initial depolarisation and primes the subsequent activation of POMC neurons; the slower mTORC1 activation induced by calcium influx and Sestrins dependent mechanism contributes to reinforce the neuronal activation machinery.

The anorectic effects of dietary proteins have shown to be mediated by multiple pathways ^10^. Based on our previous work and working hypothesis that brain leucine sensing is engaged in response to high protein feeding ^5,13^, we tested the role of MBH Cav3.1 expression in the feeding and metabolic responses to dietary proteins. As previously described, maintenance on a high protein diet reduces appetite and promotes weight loss ^32,33^. Strikingly, knockdown of *Cacna1g* in the MBH blunts the appetite-suppressive action of high-protein feeding. POMC-specific *Cacna1g* knock-down remarkably recapitulates this phenotype, suggesting that Cav3.1 in POMC neurons mediates the anorectic response to high-protein feeding.

Recent studies have revealed that Cav3.1 physically interact with TRPC5 to synergistically mediate leptin induced excitation of hypothalamic POMC neurons *in vitro* ^15^. TRPC5 has previously been shown to mediate the activation of MBH POMC neurons induced by various satiety signals such as leptin, insulin, serotonin, and GLP1-R signalling ^25,26,34,35^. In our RNAscope analyses, we observed a striking over 90% co-expression of *Trpc5* and *Cacna1g* among a half of MBH POMC neurons. Electrophysiological recordings revealed that TRPC5 is in fact required for leucine-induced activation of POMC neurons. Thus, we conclude that while Cav3.1 serves as a leucine sensor, the interaction with TRPC5 is needed to amplify the calcium flux and subsequent activation of POMC neurons following leucine exposure. This is consistent with the properties of T-type calcium channels, characterised by their low activation threshold and important in setting the excitability of neurons ^27^. The physical interaction of Cav3.1 with TRPC5 uniquely posits Cav3.1 to regulate the sensitivity of POMC neurons to other satiety signals. Consistently, we found that the Cav3.1 activator SAK3 potentiates the appetite-suppressive effect of weight-loss drugs signalling in POMC neurons via TRPC5. Together, our data suggest Cav3.1 plays a crucial role in tuning the excitability of POMC neurons in maintaining energy homeostasis.

Importantly, in a recent transcriptomics study of Prader–Willi syndrome (PWS) patients-a genetic disorder characterised by overeating and obesity – the expression of hypothalamic *CACNA1G* was found downregulated by ∼60% ^36^. These results suggest that the role of Cav3.1 in appetite control might be conserved in humans, in line with our observation that Cav3.1 is required for leucine-induced activation of human iPSC-derived POMC neurons. Here, we showed that high fat feeding downregulates MBH *Cacna1g* expression, supporting a role for decreased Cav3.1 activity in the pathophysiology of hyperphagia. Thus, deficiency of hypothalamic Cav3.1 activity might be a common pathogenic mechanism contributing to overeating in diet-induced and genetic forms of obesity. This prompted us to test whether pharmacological activation of MBH Cav3.1 might be a viable strategy to treat hyperphagic obesity. Previous work had implicated Cav3.1 in body weight control, showing that genetic and pharmacological inhibition of Cav3.1 activity in mice is protective against diet-induced obesity, independently of energy intake and specifically through increased energy expenditure ^22^. In contrast, here we found that high-fat feeding produced the same rate of weight gain in Cacna1g^MBH^ ^KO^ mice as in controls. Reduced MBH *Cacna1g* expression in response to high-fat feeding in control mice might explain the lack of significant differential phenotypes under these conditions. Nevertheless, these data argue for a role of other mechanisms in the weight-protective effect of systemic pharmacological Cav3.1 inhibition, perhaps through peripheral targets regulating energy expenditure. In the periphery, *Cacna1g* is expressed in secretory cells such as enteroendocrine cells, adrenal cortical and chromaffin cells (Human Protein Atlas). Altered secretion from these endocrine cells can drastically affect metabolic rate. Thus, we developed strategies to target Cav3.1 activity in a brain-specific manner. As expected, direct MBH injection of SAK3 suppresses food intake and potentiates the anorectic effects of other weight loss drugs. Critically, we demonstrated the feasibility of targeting MBH Cav3.1 to treat DIO via intranasal administration of SAK3. This route of central administration is highly preferred for its non-invasiveness and the degree of control for dosing time and frequency. Non-polar small molecule drugs like SAK3 are particularly suitable for this delivery route. In addition, the MBH is an ideal target as it is accessed by multiple diffusion pathways ^37^. The recent development and approval of using GLP1R agonism to treat obesity has been a major success in the face of global rise of obesity incidence. However, it has been reported that around 15-20% patients failed to response to the treatment ^38^. We showed IN delivery of SAK3 can enhance the efficacy of GLP1R agonism. Thus, IN delivery of Cav3.1 activator might benefit those patients with limited weight loss in response to GLP1R agonists. Finally, the enthusiasm for adopting high protein feeding as a long-term body weight management strategy has been limited by mounting evidence showing the associations between high circulating levels of BCAA and obesity, insulin resistance ^39–42^ and increased cardiovascular risk in both humans and rodent models ^43,44^. IN delivery of Cav3.1 activator represents a novel strategy harnessing the satiety pathways targeted by dietary proteins to manage body weight with minimal side effects.

Altogether, this study expands the current understandings of the molecular biology of cellular amino acid sensing and the neurocircuitry mediating the feeding effects of dietary protein sensing. The application of these knowledge has led to establishing a potential novel anti-obesity treatment.

## Acknowledgments

We would like to thank Prof. Pernille B. Laerkegaard Hansen (University of Southern Denmark and AstraZeneca) for kindly gifting us the fixed brains of Cav3.1 global knock out mice. At the Cancer Research UK Cambridge Institute, we would also like to thank Julia Jones (Histopathology and ISH Core Facility) and Heather Zecchini (Light microscopy Core Facility) for their assistance with RNAscope staining and imaging; and Richard Houghton and Maria Pawula for their assistance with mouse plasma amino acid analyses. We thank the Histopathology, Peptidomic, Imaging and Disease Model Cores and the Core Biochemistry Assay Laboratory at the Wellcome-MRC Institute of Metabolic Science Metabolic Research Laboratories for their help with this work. This work was supported by the Medical Research Council (MR/S011552/1; C.B. and MRC.MC.UU.00039/03; F.M.G./F.R.), the Medical Research Council Metabolic Disease Unit and Mouse Biochemistry Laboratory Grants (MC_UU_00014/5) and (MRC_MC_UU_12012/5) and the Wellcome Trust (Strategic Award 208363/Z/17/Z for the MRL Disease Model Core Facility, Genomics and Transcriptomics Core Facility, Proteomics and Peptidomics Core Facility and Imaging Core Facilities and 220271/Z/20/Z; F.M.G./F.R.). F.T.M. is a New York Stem Cell Foundation - Robertson Investigator [NYSCF-R-156] and Chan Zuckerberg Initiative - Ben Barres Early Career Investigator [CZI NDCN 191942, 10.37921/429861umrcjh], and is supported by the Wellcome Trust and Royal Society [211221/Z/18/Z]. For the purposes of open access, the authors have applied a CC-BY public copyright license to any Author Accepted Manuscript version arising from this submission.

## Author Contributions

C.B. conceived the study and acquired fundings. A.H.T., N.H. C.A.A., E.H., B.L., T.D., D.N., R.G.A., A.S., R.W., N.B., A.P., C.B. researched and analysed data. T.F. performed in silico docking analysis and provided expert opinions. A.H.T., N.H., C.A.A. E.H. B.L., G.S.Y., F.T.M., F.M.G., F.R., K.W. and C.B. designed experiments and wrote and edited the manuscript. G.S.Y., F.T.M., F.M.G., F.R., K.W. and C.B. supervised the projects. P.K. and M.M. provided crucial reagents and assistance. All authors reviewed and edited the manuscript.

## STAR Methods

Details of all reagents and animal models are detailed in the **Key Resource Table**.

### Resource availability

#### Lead contact

Further information and requests for reagents may be directed to and will be fulfilled by the Lead contact, Clemence Blouet

#### Materials availability

This study generated following unique reagents: 1.) various stable cell lines and plasmids of hCav3.1 mutants, and 2.) two new anti-mCacna1g gRNA AAV plasmids which are available upon request.

#### Data and code availability

Sequencing data of PhosphoTRAP are available on GEO (number pending). All other data to understand and assess the conclusion of this research are available in the main text and supplementary materials.

### Experimental model and Subject details Mouse strains and husbandry

All animal experiments (except for the mouse brain slice electrophysiology experiments performed at the University of Texas as noted below) were performed in accordance with the UK Home Office regulations under the Animals (Scientific Procedures) Act (1986) and with the approval of the University of Cambridge Animal Welfare and Ethics Review Board.

Wild-type C57BL/6J mice were obtained from Charles River UK (at 7 weeks of age) and were allowed to acclimatise to the facility for 1 week prior any experimental procedures. The following transgenic mouse strains were obtained from the Jackson Laboratory: P*omc-eGFP* (Tg(Pomc-EGFP)1Low/J; Strain #:009593) ^45^, *Pomc-Cre* (Tg(Pomc1-cre)16Lowl; Strain #:005965) ^46^, *R26-LSL-Cas9::EFGP* (tm1(CAG-cas9*,-EGFP)Fezh; Strain #:026175) ^47^. All transgenic mouse lines were kept on C57BL/6J genetic background. For all the experiments using Cre reporter lines, we conducted the work in hemizygous males or wild-type littermates randomly assigned to the experimental groups. For studies on wild-type mice, weight-matched groups were compared.

Mice were group-housed in individually ventilated cages with standard bedding and enrichment, and maintained in a humidity-controlled room at 22–24°C on a 12-hr light/dark cycle with ad libitum access to water and standard laboratory chow diet (SAFE® 105, SAFE® Complete Care Competence, France) unless otherwise stated. Isocaloric modified diets with varying protein amounts were custom made by Research Diets as per the formulations in Table S6. 60% high fat diet was obtained from Research Diets (Cat# D12492). Before each dietary change, the mice were briefly exposed to the new diets to avoid neophobia or other novelty-related responses in subsequent experiments. Mice were acclimatised to the dosing procedures for at least 4 consecutive days prior any dosing studies.

For brain slice electrophysiology experiments performed at the University of Texas, male (6- to 18-week-old) pathogen-free mice were used for all experiments. All mice were housed under standard laboratory conditions (12 hr light-dark cycle) and a temperature-controlled environment with food and water available ad libitum. All experiments were performed in accordance with the guidelines established by the National Institute of Health Guide for the Care and Use of Laboratory Animals and approved by the University of Texas Institutional Animal Care and Use Committee. To identify POMC neurons, *Pomc-hrGFP* (Tg(Pomc1-hrGFP)1Lowl/J, Strain #:006421) mice were utilized ^47^. *Trpc5* KO (Trpc5tm1.1Clph) mice ^48^ were subsequently mated with *POMC-hrGFP* to identify POMC neuron on a *Trpc5* KO background.

### Primary culture of post-weaning mediobasal hypothalamic neurons

Acute primary cultures of mediobasal hypothalamic neurons were prepared from 5 to 6-week-old POMC-EFGP male mice as previously described ^6^. Briefly, mice were fasted overnight and sacrificed with cervical dislocation. Brains were extracted into an ice-cold custom-formulated extraction media based on rat cerebrospinal fluid and microdialysis extracts ^5^. For each brain, 2 brain sections of 0.75 mm were cut using a McIlwain tissue chopper, starting 1.2 mm post the anterior commissure and ranging from −1.06 mm post Bregma to −2.54 mm post Bregma. The MBH was then dissected from coronal slices using a dissecting microscope. Tissue was transferred to papain (20 U/ml, Worthington, Lakewood, NJ, USA) pre-heated at 37°C and digested for 30 min at 37°C under agitation (Thermomixer, 500 rpm). After digestion, tissue extracts from 6 to 7 animals were pooled, transferred to a tube containing extraction media with 3.5 U/ml DNase I from bovine pancreas (Sigma) using a glass Pasteur pipette with a fire polished 1.5 mm opening and triturated with pipettes. The trituration supernatant was gently loaded on top of a BSA gradient (4% BSA prepared in extraction media (pH 7.4) loaded on top of 8% BSA (pH 7.4)), centrifuged for 5 min at 300 rcf, and the pellet was resuspended in a custom-formulated culture media (ref). 100 μl of resuspended cells were plated on the poly-D-lysine coated glass-bottom 35 mm dishes (MatTek Corporation) using a 0.5 mm trituration pipette, inside a cloning cylinder (8 mm^2^, Sigma). Plates were placed in an incubator (37°C, 5% CO2) for 1 hr. After that, an additional 2 ml culture media was added and the cloning cylinder removed. 4 to 6 culture dishes were prepared on each experimental day. Each culture dish was imaged once and represented our experimental unit.

### Human derived hypothalamic neurons

Human induced pluripotent stem cells (FSPS13B line) were maintained in mTESR1 media and differentiated to hypothalamic neurons as previously described ^49^. Human hypothalamic neurons were allowed to differentiated for 30–50 days in vitro, at which point there were abundant neurons immunoreactive for POMC ^49,50^. To study cell-autonomous responsiveness to leucine, cultures were enzymatically dissociated with TrypLE and Papain as described previously (ref) and replated onto glass-bottomed 35 mm dishes (MatTek Corporation) coated with Geltrex (Thermo Fisher Scientific, 21041025). Cells were loaded with 5 μM Fura2 AM for 45 min and treated with vehicle or experimental solutions.

### Brain slice preparation

Brain slices were prepared from young adult male mice (6-10 week-old) as previously described ^25,35,51^. Briefly, male mice were deeply anesthetized with i.p. injection of 7% chloral hydrate and transcardially perfused with a modified ice-cold ACSF (described below). The mice were then decapitated, and the entire brain was removed and immediately submerged in ice-cold, carbogen-saturated (95% O2 and 5% CO2) ACSF (126 mM NaCl, 2.8 mM KCl, 1.2 mM MgCl2, 2.5 mM CaCl2, 1.25 mM NaH2PO4, 26 mM NaHCO3, and 5 mM glucose). Coronal sections (250 mm) were cut with a Leica VT1000S Vibratome and then incubated in oxygenated ACSF (32°C-34°C) for at least 1 hr before recording. The slices were bathed in oxygenated ACSF (32°C-34°C) at a flow rate of ∼2 ml/ min. The pH and osmolality of all ACSF with or without L-Leucine adjusted to within 7.2-7.3 and 290-300 mOsm/L range respectively. All electrophysiology recordings were performed at room temperature.

### Cell culture

Human embryonic kidney HEK293 cell line (ATCC) and mouse embryonic hypothalamic mHypoE-N46 cell line (Cedarlane Labs, NC, USA) were maintained in DMEM with 2 mM stable glutamine supplemented with 10% fetal bovine serum (FBS) and 10,000 U penicillin/streptomycin at 37°C with 5% CO_2_. The cells were authenticated by STR profiling (Eurofins) and tested negative for mycoplasms by PCR (Merck) regularly. Cells were used for experiments before 30 passages. Lipofectamine 3000 (Thermo Fisher Scientific) were used for transient transfection of mHypoE-N46 cells according to manufacturer’s protocol (Publication No. MAN0009872 Rev C.0). TransIT-293 Transfection Reagent (Mirus Bio) was used for transfecting HEK293 cells according to manufacturer’s protocol (ML009-Rev.G 0117).

HEK293 cells stably expressing hCav3.1 (HEK293-hCav3.1) were generated monoclonally. In brief, 48hr after transient transfection with linearised hα1Ga-pDsRed plasmid (Addgene #45811) ^52^, cells were selected with antibiotic G418 at 750 μg/ml for about 2 weeks. Single clones of resistant cells were generated with serial dilutions and retrieved with the aid of cloning rings. This stable cell line was maintained with standard medium supplemented with 250 μg/ml G418 and was used for all cell experiments involving hCav3.1 except those involved truncated hCav3.1 and site-specific mutated hCav3.1.

HEK293 cells stabling expressing hCav3.1 site-specific mutations and their wildtype control (WT) were generated polyclonally. 48 hr after transient transfection with the linearised hCav3.1 WT and mutant plasmids with puromycin-resistant gene, cells were selected with antibiotic puromycin at 2 μg/ml for about 2 weeks. The resultant stable cell lines were maintained with standard medium supplemented with 1 μg/ml puromycin and was used for cell experiments involving site-specific mutations of hCav3.1.

## Method details

### Stereotaxic surgical procedures and adeno-associated virus (AAV) vectors

Surgical procedures were conducted on 9- to 11-week-old mice under isoflurane anaesthesia. All the animals received Metacam prior to- and 24 hr post-surgery, and were allowed a 1-week recovery period during which they were acclimatized to injection procedures. For brain viral injections under anaesthesia, mice were stereotactically implanted with bilateral steel guide cannulas (Plastics One) positioned 1 mm above the ARH (A/P: 1.1 mm, D/V: 4.9 mm, and lateral: +/−0.4 mm from the bregma). Beveled stainless steel injectors (33 gauge) extending 1 mm from the tip of the guide were used for injections. For chronic cannula implantation, the cannula guide was secured in place with Loctite glue and dental cement (Fujicem2). Correct targeting was confirmed histologically post-mortem. The mice were allowed 1 week of recovery during which they were handled daily and acclimatized to the relevant experimental settings. For AAV injections, 500 nl AAVs per side were injected into the ARH with an injection rate of 100nl per minute and were kept in place for additional 5 minutes post-injection under anaesthesia. After that, the wound was closed by surgical suturing and allowed for 3 weeks recovery and transgene expression. The following AAVs produced by Vector Biolabs were used in this study: AAV/DJ-U6-anti-Cacna1g gRNA-CMV-SaCas9 (5.2 × 10^12^ vg/ml); AAV/DJ-CMV-SaCas9 (6.3 × 10^12^ vg/ml); AAV/DJ-U6-dual anti-Cacna1g gRNA-hSyn-mCherry (4.7 × 10^12^ vg/ml); AAV/DJ-hSyn-mCherry (7.1 × 10^12^ vg/ml).

### MBH injection on behaving mice and food intake assessments

The same general design applied across all experiments involving MBH injection: studies were conducted in a home-cage environment; mice were food-deprived for 6hr during the day-time before receiving a bilateral parenchymal injection (100 nl/side, and 50 nl/min) of following drugs: L-leucine 2.1 mM (Sigma)’ SAK3 0.1-1mM (Tocris), Leptin 5 μg/μl (R&D Systems), liraglutide 1 μg/μl (Tocris), lorcaserin 10 μg/μl (Generon) or control aCSF with matching DMSO content as a vehicle. Mice were immediately returned to their home cage for food intake analysis or culled for specific experiments as per indicated time intervals.

Mice were fasted from ZT5-11 for PhosphoTRAP assay and from ZT3-9 for RNAscope in situ hybridization and immunofluorescence histological assessments. They were culled with specific details in respective sections below.

For food intake studies, mice were fasted at ZT5 and the injection occurred 1 hr before dark onset. The mice were refed after the injection and their food intake was monitored over various time points after refeeding. All the studies were conducted in a cross-over randomized manner on age- and weight-matched groups, and at least 4 days elapsed between each brain injection.

### PhosphoTRAP assay

To identify enriched transcripts in mediobasal hypothalamic neurons activated or inhibited by the administration leucine compared to aCSF, PhosphoTRAP assay was performed with adaptions as previously described ^9,53^. 45 min after the MBH injection as described above, mice were culled by overdose of anaesthetic (300 mg/kg I.P. sodium pentobarbitone; Euthatal Solution for Injection, Dopharma Research B.V.). The MBH was micro-dissected in an ice-cold Buffer B (Hanks Balanced Salt Solution (HBSS) with 2.5 mM of HEPES at pH 7.4, 4 mM of NaHCO3, 35 mM of glucose, and 100 mg/mL of cycloheximide in methanol) under a 10x dissecting microscope. The hypothalami of 5 mice were pooled as a single sample resulting in 4 replicates per experimental condition (i.e. aCSF vs. Leu). Pooled hypothalami were manually homogenized in a 1ml of buffer C (10 mM of HEPES at pH 7.4, 150 mM of KCl, 5 mM of MgCl2, 100 nM of calyculin A, 2 mM of DTT, 100 U/mL of RNasin, 100 mg/mL of cycloheximide, and Roche protease and phosphatase inhibitor cocktails) and clarified by centrifugation at 2,000 g at 4°C for 10 min. The supernatants were transferred to a new tube. 70μl of 10% NP40 and 70μl of DHPC (1,2-diheptanoyl-sn-glycero-3-phosphocholine) were added, the samples were then mixed by inversion and allowed for 2 min incubation on ice, and centrifuged at 16,100 g at 4°C for 10 min. Supernatants were transferred to a new tube. 25 μl of each sample was collected and subject to RNA extraction as “Input sample”. The remaining sample was used for ribosome immunoprecipitation (IP).

Prior to the immunoprecipitation step, 100μl Protein A Dynabeads (Invitrogen) were washed 3 times with buffer A (10mM HEPES, 150mM KCl, 5mM MgCl2 and 1% NP40 (pH 7.4) before incubation with 4μg pS6 antibody against Ser240/244 (#2215, Cell Signalling Technology) and 0.1% bovine serum albumin (BSA) in buffer A (300 μl per IP sample) at 4°C overnight with continuous mixing on an end-over-end rotator. On the next day, the antibody-bead conjugates were washed twice with wash buffer D (10 mM of HEPES, 350 mM of KCl, 5 mM of MgCl2, 2 mM of DTT, 1% NP40, 100 U/mL of RNasin, 100 mg/mL of cycloheximide, and Roche protease and phosphatase inhibitor cocktails). After the last wash, the beads were resuspended in 200 μl of homogenisation buffer C supplemented with 50 μl of 10% NP40 and 10 μl of DHPC for 1 mL of buffer C and 1 μM of ZK10 (a gift from Dr. Zackary Knight, UCSF, US). The remaining sample supernatant was added to the antibody-beads conjugates, resuspended by pipetting and mixed with a rotator at 4°C for 10 min. The beads were washed 4 times with 0.9 mL of ice-cold wash buffer D and resuspended in 350 μl of RLT buffer and allowed for 5 min incubation on ice. The supernatant was then collected as “IP sample.”

RNA was extracted using the RNeasy Micro kit (#74004; Qiagen). RNA quantity was assessed using the Quant-iT RiboGreen RNA Assay kit (#R11490; Thermo Fisher Scientific), and the samples’ RNA quality was assessed using Pico chips on an Agilent Bioanalyser. The samples were cleared from DNA contamination using a TURBO DNA-free kit (#AM1907; Thermo Fisher Scientific). cDNA was prepared using a Smart Seq v4 Ultra Low Input RNA kit (#634889; Clontech). Library preparation was done using a Nextera XT kit (Illumina), and the samples were run on a NexSeq 500 System (#FC-131-1024; Illumina).

### PhosphoTRAP pathway analysis

Ingenuity Pathway Analysis (QIAGEN) was used to identify pathways that are up- or downregulated in each cluster between experimental conditions. Log2FC values, p values, and FDRs for each gene for each cluster were uploaded to the software.

### Metabolic phenotyping

Body composition was analysed using a EchoMRI Whole Body Composition Analyser (EchoMRI, Houston, Texas). Promethion High-Definition Multiplexed Respirometry Cages (Sable Systems International, Las Vegas, Nevada) were used to analyse energy expenditure by indirect calorimetry, food intake, water intake, respiratory quotient and locomotor activity over 48 hr in mice single-housed for at least one week prior. Data collected during the first 24 hr of each run was discarded to allow for acclimatisation of mice to the altered cage environment.

### Glucose phenotyping

For the oral glucose tolerance test, mice were food deprived for 6 hr from ZT1-7 and blood was sampled from the tail vein immediately prior to glucose bolus (gavage, 2 mg/kg), and 10, 20, 30, 60, 90, and 120 min following bolus administration. For the insulin sensitivity test, mice were food deprived for 6 hr from ZT1-7 and blood was sampled from the tail vein immediately prior to the administration of an insulin bolus (i.p., 0.75 U/kg), and 10, 20, 30, 60, 90, and 120 min following bolus administration. Blood glucose was analysed using an AlphaTrak3 handheld glucometer (Precision Xtra; MediSense).

### Intranasal administration of substances on awake mice

Intranasal (IN) administration utilising a modified scuffing method to immobilise awake mice during drug administration via the nostrils was performed according to an established protocol ^54^. Briefly, mice were acclimatised for the modified scuffing method and mock injection (air-puffing only from pipette) either daily for two weeks in the chow-fed pilot experiment or bi-weekly over 12 weeks in the high fat diet experiment. On the day of actual experiment, a total of 24 μl of solution was applied to both nostrils of a mouse, over 4 sessions of 6 μl administration, ejecting slowly from a gel-loading pipette tip to allow formation of small droplets and inhalation by the mouse.

For pilot experiments, the following drugs were IN administered 1 hr before dark onset: 1 mM SAK3, 1 μg/ μl liraglutide or vehicle control (1% DMSO in saline) content. Food and mice were weighed before and 24 hr after the IN administration. Each of the testing conditions was separated by 4 days apart.

For the experiment on HF-fed mice, 7 week-old WT mice were transited to and maintained on 60% high fat diet for 12 weeks; upon then, only mice weighing over 40 g were used. Mice first received daily IN administration of vehicle (1% DMSO in saline) and subcutaneous (SC) injections of saline (5 ml/kg) during ZT 10-11 for 2 weeks to obtain an individual baseline response to the procedures and then daily I.N. administration of saline. After that, mice were proceeded to receive I.N. administration of 1mM SAK3 (or vehicle) with SC injection of 0.2 mg/kg liraglutide (or vehicle) during the same time-of-day for another 2 weeks. Food and animals were weighed daily throughout the study.

### Brain tissue preparation for histological assessments

Mice were anaesthetised with an intraperitoneal injection of 100 mg/kg sodium pentobarbitone (Dolethal, Vetoquinol UK Ltd) then transcardially perfused with chilled 0.1 M heparinized phosphate buffered saline (PBS) followed by 4% paraformaldehyde (pH 7.4). The brains dissected and post-fixed overnight at 4°C in 4% PFA then cryoprotected in 30% (w/v) sucrose solution in PBS with 0.03% sodium azide as a preservative for at least 48 hr prior to processing. For RNA-scope experiments, the cryoprotected brains were embedded in Optimal Cutting Temperature (OCT) medium (CellPath, Newtown, UK) in plastic molds, frozen on crushed dry ice and then stored at −80°C until further processing.

### Brain tissues preparation for simultaneous gDNA, RNA and protein isolation

To validate the CRISPR mediated MBH Cacna1g KO models, the micro-dissected mediobasal hypothalami of control and Cacna1g^MBH^ ^KO^ mice were collected in the RNAlater solution (Thermo Fisher Scientific) and stored according to the manufacturer’s protocol (7020M Rev. G 15Jul2014). On the day of processing, the tissues were removed from RNAlater solution and then homologized with a hand-held electric homogeniser and a pistil (Sigma Aldrich) in 300 μl TRIzol Reagent (Thermo Fisher Scientific) to simultaneously isolate of gDNA, total RNA and protein according to the manufacturer’s protocols (MAN0001271 Rev. B.0 and MAN0016385 Rev. A.0).

### Multiplexed Fluorescent in situ hybridization (FISH) with RNAscope

Simultaneous detection of mouse *cFos, Trpc5, Cacna1g and Pomc* was performed on frozen sections using Advanced Cell Diagnostics (ACD) RNAscope 2.5 LS Multiplex Reagent Kit (Cat No. 322800), RNAscope 2.5 LS Probe Mm-Fos (Cat No. 316928), RNAscope 2.5 LS Probe Mm-Trpc5 (Cat No. 476248), RNAscope® 2.5 LS Probe Mm-Cacna1g-C2 (Cat No. 459768 C2), RNAscope® 2.5 LS Probe Mm-POMC-C3 (Cat No. 314088-C3), (ACD, Hayward, CA, USA). Briefly, OCT embedded frozen brains were sliced at 12 µM using a Leica CM1950 cryostat directly onto Superfrost Plus slides (Thermo Fisher Scientific) in an RNase-free environment and stored at −80°C until use. Upon processing, slides were thawed at room temperature for 10 min and post-fixed in pre-chilled 4% PFA for 15 min at 4oC, washed in 3 changes of PBS for 5 min each before dehydration through 50%, 70& 100% Ethanol for 5 min each. The slides were air-dried for 5 min before loading onto a Bond Rx instrument (Leica Biosystems). Slides were prepared using the frozen slide delay prior to pre-treatments using Epitope Retrieval Solution 2 (Cat No. AR9640, Leica Biosystems) at 95°C for 2 min, and ACD Enzyme from the Multiplex Reagent kit at 40°C for 15 min. Probe hybridisation and signal amplification was performed according to manufacturer’s instructions.

Opal 650 (Akoya Biosciences Cat No. FP1496A) detection at 1:750 dilution of Trpc5 or Fos, Opal 570 (Akoya Biosciences Cat No. OPI-001003) detection at 1:750 of Cacna1g, and Opal 690 (Akoya Biosciences Cat No. OPI-001006) detection at 1:3000 dilution of POMC were performed on the Bond Rx according to the ACD protocol. Slides were then removed from the Bond Rx and mounted using Prolong Diamond (Thermo Fisher Scientific).

The brain sections were imaged using a spinning disk Operetta CLS (PerkinElmer) in confocal mode using an sCMOS camera and a 40x automated water-dispensing objective. The sections were imaged with z stacks at intervals of 1 μm. ROIs included the ARH and VMH. Gain and laser power settings remained the same between the experimental and control conditions during each experiment. Harmony software (PerkinElmer) was used to automatically quantify the number of labelled RNA molecules (spots) per cell and the number of labelled cells among other metrics.

### Immunohistochemistry and imaging

Perfused brains were post-fixed overnight at 4°C in 4% PFA then cryoprotected in 30% (w/v) sucrose (Fisher Scientific) solution in PBS for at least 48 hr prior to processing. Tissues were covered with Optimal Cutting Temperature (OCT) medium (CellPath, Newtown, UK) and sections were obtained at 25 μm on a Leica SM2010R Freezing Microtome (Leica, Wetzlar, Germany). All sections were subjected to heat-mediated antigen retrieval in 10 mM sodium citrate (pH6.5; Fisher Scientific) in distilled water for 20 min at 80°C prior to washing 3 times in PBS. For all experiments, sections were blocked in normal goat serum (NGS, Abcam) diluted in PBS containing 0.3% Triton X-100 (0.3% PBST; Sigma) for 1 hr prior to primary antibody incubation overnight at 4°C. Following primary antibody incubation, sections were washed 3 time with 0.1% PBST and incubated with appropriate fluorophore-conjugated secondary antibodies diluted 1:500 in 0.3% PBST for 2 hr at room temperature. Sections were subsequently washed with 0.1% PBST and mounted to Clarity microscope slides (Dixon Science, Edenbridge, UK) under coverslips (1.0 thickness; Marienfeld, Lauda-Königshofen, Germany) with Vectashield Vibrance Mounting Medium with 4ʹ,6-diamidino-2-phenylindole (DAPI; Vector Laboratories, Newark, California). The following primary antibodies were used: anti-Phospho-S6 Ribosomal Protein (Ser240/244) (D68F8) Rabbit monoclonal-Alexa Fluor 594 Conjugate) (1:200; Cell Signalling Technology, Cat# 9468), anti-cFOS rabbit monoclonal and guinea pig monoclonal (1:1000; Synaptic Systems, Cat# 226008 and 226308, respectively), anti-Cav3.1 (C-terminal) rabbit polyclonal (1:200; Abcam, Cat# ab203577), anti-GFP chicken polyclonal (1:1000; Abcam, Cat# ab13970), anti-DsRed mouse monoclonal (1:1000, Clontech, Cat# 632392), anti-POMC rabbit polyclonal (1:1000; Phoenix Pharmaceutical, Cat# H-029-30).

The sections were imaged using a Zeiss Axio slide scanner with a 20x objective or a Leica SP8 confocal microscope with 40x or 63x objectives. The imaging settings remained the same between the experimental and control conditions. Images of tissue sections were digitized, and areas of interest were outlined based on cellular morphology and using Paxinos and Franklin’s brain atlas [31]. The images were analyzed using the ImageJ manual cell counter or Zeiss ZEN 2.3 software.

### Quantitative real-time polymerase chain reaction (qPCR)

Total RNA of tissues and cell cultures was extracted using either TRIzol reagent according to the manufacturer’s protocol (MAN0001271 Rev. B.0). cDNA synthesis of 1 μg RNA was performed using the High-Capacity cDNA Reverse Transcription Kit with random hexamer primers (Thermo Fisher Scientific). qPCR was performed using either TaqMan Universal PCR Master Mix for reaction based on dual-labelled probes or the SYBR Green PCR Master Mix for SYBR Green reactions (both from Thermo Fisher Scientific) on a QuantStudio 5 Real-Time PCR System (Thermo Fisher Scientific). The average of technical duplicates of a biological sample was used for quantification. Relative gene expression was quantified using the ΔΔ threshold cycle (Ct) method with adjustments to the amplification efficiencies of individual primer pairs. *Eef1α* was used as the reference gene for all experiments. The specificity of all SYBR Green primers were validated with melting curve analyses. All custom-designed standard qPCR primers and 6-FAM-TAMRA dual labelled probes were designed to target the exon-exon boundary of a gene or separated by a long intron to exclude amplifying genomic DNA, and were synthesised by Merck. Predesigned dual-labelled TaqMan Gene Expression Assay probes were purchased from Thermo Fisher Scientific. Information for oligos used are shown in the **Key Resources Table**.

### SDS-polyacrylamide gel electrophoresis (PAGE) and Western blot

Protein from MBH tissues were extracted with TRIzol Reagent as described above. Protein from cultured cells were lysed with RIPA lysis buffer (1% triton X-100, 2% sodium dodecylsulfate (SDS), 1% sodium deoxycholate, 1 % NP40 and 1x cOmplete protease inhibitor cocktail (Roche, Grenzach-Wyhlen, DE) in Tris-buffered saline (TBS) and subjected to sonication using a Branson 450 sonifier (Thermo Scientific; amplitude: 50, duty: 30%, duration: 30 s). In the case of detecting phosphorylated proteins, 1x Roche pSTOP phosphatase inhibitors was included to the lysis buffer. Protein concentrations were determined by BCA protein assay kit (Thermo Scientific). Appropriate protein amount was further diluted with SDS sample-loading buffer (50mM Tris-HCl pH 6.8, 0.05% (w/v) bromophenol blue, 6.7% glycerol, 1% SDS, 1.5% β-mercaptoethanol).

Due to the large molecular mass of Cav3.1 (∼262 kDa), a modified immunoblotting protocol was used for detecting Cav3.1. For detecting Cav3.1, samples were heated at 70°C for 10 min. 60 μg protein from MBH tissues or 40 μg from cultured cells were loaded per sample onto a 4 – 12% pre-casted Novex Tris-Glycine Mini Protein Gels (Thermo Fisher Scientific). After electrophoresis, gel was first equilibrated in a 2x Tris-Glycine Transfer buffer (25 mM Tris, 192 mM glycine, pH 8.3) containing 0.04% SDS for 10 min. After that, Western blotting was performed on a methanol-activated PVDF membrane using 1x Tris-Glycine Transfer buffer containing 10% methanol and 0.01% SDS at 12 V overnight or 20 V for 3h. Both electrophoresis and Western blotting were carried out with the Invitrogen Mini Blot system (Thermo Fisher Scientific).

For detecting other proteins, samples were heated at 95°C for 5 min. 40 μg protein from cell culture were loaded per sample. A standard SDS-PAGE and Western blotting protocol was used as described above except the following differences: 1.) standard 10% PAGE gel was used; 2.) the pre-equilibration step was omitted; 3.) the Transfer buffer contained 20% methanol and no SDS; 4.) The transfer was carried out at 20V for 2 hr. 5% non-fat milk (Bio-Rad) was used as the blocking agent for primary antibodies detecting non-phosphorylated proteins and all horseradish peroxidase (HRP)-conjugated secondary antibodies (Abcam, Cat# ab205718-ab205719, respectively). 5% BSA was used as the blocking agent for primary antibodies detecting phosphorylated proteins. SuperSignal West Pico PLUS Chemiluminescent Substrate (Thermo Fisher Scientific) was used for developing the chemiluminescence signal of the blot, which was then imaged and digitalised with the ChemiDoc Imaging System (Bio-Rad). Densitometric analysis of band intensity was performed with Quantity One software (Bio-Rad). Quantification across multiple blots were normalised with a shared calibrating samples with the same amount of protein. Endogenous GAPDH on the same blot was used as the internal loading control for standard Western blot experiments. Total non-phosphorylated form of the target proteins from the same blot was used as the internal loading control for phosphorylated protein after stripping with the Restore PLUS Western Blot Stripping Buffer (Thermo Fisher Scientific).

The following primary antibodies were used: anti-Cav3.1 (C-terminal) mouse monoclonal (1:1000; abcam, Cat# ab134269), anti-TRPC5 rabbit polyclonal (1:1000; Alomone Labs, Cat# ACC-020), anti-GAPDH mouse monoclonal (1:1000; Santa Cruz Biotechnology, Cat# sc-32233), anti-phospho-p70 S6 Kinase (Thr389) rabbit polyclonal (1:1000; Cell Signalling Technology, Cat# 9205), anti-P70 S6K rabbit monoclonal (1:1000; Cell Signalling Technology, Cat# 2708) and anti-HA mouse monoclonal (1:1000; Cell Signalling Technology, Cat# 2367).

### Blood amino acid profiling

Whole blood was collected terminally into a EDTA-treated tube (Sarstedt) using right ventricle cardiac puncture from mice under anaesthesia following an intraperitoneal injection of 100 mg/kg sodium pentobarbitone. Plasma was retrieved as supernatant after centrifuging the whole blood at 4°C at 2000 g for 15 mins. 5 µl of plasma sample was added with mixed isotope-labelled amino acid internal standards, then extracted by protein precipitation using formic acid and isopropyl alcohol followed by derivatisation using AccQ-Tag Ultra Derivatising Kit (Water 186003836). Derivatized amino acid standards in 0.1% formic acid was prepared and analysed along with the extracted samples. Liquid chromatography and tandem mass spectrometry was performed on the Shimadzu Nexera X2 Liquid chromatography system linked to Sciex 6600 Triple Quad mass spectrometer. 19 proteinogenic amino acids (except glycine due to protocol limitations) plus citrulline, orthenine and taurine were analysed.

### Urine analyses

Urine was collected from mice during ZT8-10 after at least 3 weeks on indicated diets on two separated days without other procedures with gentle abdominal massaging to encourage urination on a sheet of water repulsive parafilm. Urinary albumin was analysed with a mouse albumin ELIZA kit (Cat# E99-134, Bethyl Laboratories, US); creatinine was measured with the Creatinine Assay Kit (Cat# EXO-1012, Ethos Biosciences, US); urea was measured with urea assay kit (Cat# ab234052, Abcam) and glucose was measured with glucose assay (Cat# ab65333, Abcam). Urinary osmolarity was measured by the Osmomat 3000 osmometer (Gontec, DE).

### SAK3 LC-MS/MS analysis

Tissues were weighed and transferred into vials containing Lysing Matrix D beads (MP Bio) along with 500 µl of 80% ACN in water. Samples were homogenised on a VelociRuptor (SLS) homogenisation system at 4.5 m/s for 1 min and were cooled on ice between each of the three cycles. The heart tissues required a second homogenisation step with the addition of a metal ball to break apart the tough muscle tissue. Samples were centrifuged for 10 minutes at 12000 g and 250 µl of the supernatants was transferred to an Eppendorf LoBind plate. The supernatants were evaporated under oxygen free nitrogen at 40 °C and reconstituted into 100 µl of 10% ACN in water and 50 µl of internal standard (D6 memantine) was added. A calibration line of SAK3 in 0.1% formic acid was prepared and analysed along with the extracted samples. Liquid chromatography and tandem mass spectrometry was performed on a Waters M-class Acquity linked to a TQ-XS mass spectrometer. A selected reaction monitoring (SRM) method was used targeting SAK3 (369.99/323.92) and labelled Memantine (180.2/163.04). The data was processed on TargetLynx XS (v 4.2, Waters) and SAK3 content was normalized by tissue weight.

### Calcium imaging

Cells were loaded with 5 μM Fura2 AM dye (Life Technologies) for 30 min, washed with custom-formulated extraction media based on rat cerebrospinal fluid ^6^, and imaged using an inverted fluorescence microscope (Olympus IX71, Olympus, Southend on Sea, UK) with a 40× oil-immersion objective lens. GFP (to identify POMC cells) was excited at 488 nm and fura-2 at 340 nm and 380 nm using a monochromator (Cairn Research, Faversham, UK) and a 75 W xenon arc lamp, and emissions were recorded using an Orca ER camera (Hamamatsu, Welwyn Garden City, UK), a dichroic mirror and a 510 nm long pass filter. All images were collected on MetaFluor software (Molecular Devices, Wokingham, UK). The ratio of fura-2 emissions at 340 and 380 nm (340/380 ratio) was used to monitor changes in the intracellular calcium concentration. Solutions were perfused continuously at a rate of approximately 0.5 ml/min.

### Whole cell patch-clamp electrophysiological recording on brain slices

The pipette solution for whole-cell recording was modified to include an intracellular dye (Alexa Fluor 350 hydrazide dye) for whole-cell recording: 120 mM K-gluconate, 10 mM KCl, 10 mM HEPES, 5 mM EGTA, 1 mM CaCl2, 1 mM MgCl2, and 2 mM MgATP, 0.03 mM Alexa Fluor 350 hydrazide dye (pH 7.3). Epifluorescence was briefly used to target fluorescent cells, at which time the light source was switched to infrared differential interference contrast imaging to obtain the whole-cell recording (Zeiss Axioskop FS2 Plus equipped with a fixed stage and a QuantEM:512SC electron-multiplying charge-coupled device camera). Electrophysiological signals were recorded using an Axopatch 700B amplifier (Molecular Devices); low-pass filtered at 2-5 kHz, and analysed offline on a PC with pCLAMP programs (Molecular Devices). Membrane potentials and firing rates were measured from POMC neurons in brain slices. Recording electrodes had resistances of 2.5-5 MΩ when filled with the K-gluconate internal solution.

A change in membrane potential was required to be at least 2 mV in amplitude in response to drug application. Membrane potential values were not compensated to account for junction potential (−8 mV). Effects of L-Leucine on AP frequency (over 0.5 Hz) before and during acute L-Leucine bath application were analysed within a recording using the Kolmogorov-Smirnov (K-S) test (a nonparametric, distribution-free goodness-of-fit test for probability distributions).

### Whole cell patch-clamp electrophysiological recording on HEK293 cell

HEK293 cells were transfected with pCAG-hCav3.1-ires-EGFP plasmid using TransIT-293 Transfection Reagent 48-72 hours before experiments. Transfected cells were plated in 35mm dishes the day before the experiments. Recordings of single EGFP+ cells were performed at room temperature (20-24ᵒC). Regular bath solution or bath solution containing 1 mM L-Leucine (prepared fresh on day of experiment) were applied directly onto cells using a custom-made gravity fed perfusion system. A constant flow of bath solution was applied for baseline recordings. The internal pipette solution contained (in mM): 115 CsCl, 1 MgCl2, 0.9 CaCl2, 20 HEPES, 5 Mg-ATP, 11 BAPTA, Ph 7.3 with CsOH.

Bath solution contained (in mM): 10 CaCl2, 140 TEA-Cl, 10 HEPES, 10 D-glucose, 2.5 CsCl and 1 MgCl2 Ph 7.3 WITH TEAOH. Cells were recorded in conventional whole cell configuration after obtaining a gigaΩ seal and recorded using an Axopatch 200B amplifier connected through a Digidata 1440A A/D converter and pclamp software (Molecular Devices). Microelectrodes were pulled from borosilicate glass (1.5 mm OD, 1.17 mm ID; Harvard apparatus) using a PC-100 Narishige microelectrode puller (Narishige, Japan) and the tips coated with refined yellow beeswax. Electrodes were fire polished using an MF-830 micro forge (Narishige, Japan) and had resistances of 2-3 MΩ when filled with internal pipette solution. A silver/AgCl ground wire connected to the bath solution via a 3M KCl agar bridge was used as a ground. To obtain current-voltage relationships cells were challenged to a 2-step activation/inactivation protocol, with an initial holding potential of −100 mV and stepped from −90 to −10 mv, in 10 mv steps (500 ms first duration, 2 seconds step to step). I/V plots, inactivation constants and raw traces were obtained and analysed using Clampfit software (Molecular Devices).

### Plasmid construction

The guide RNAs used were designed to target an early exon common to all transcript variants of mouse *Cacna1g* with the aid of the CRISPick gRNA design online tool (Broad Institute). The selected custom gRNAs in pX601-AAV-CMV::NLS-SaCas9-NLS-3xHA-bGHpA vector (for Cacna1g^MBH^ ^KO^) ^55^ and pSpCas9(BB)-2A-GFP PX458 vector (for Cacna1g^POMC^ ^KO^) ^56^ were purchased directly from Genescript (company) and validated on mHypoE-N46 cells for CRISPR-mediated indel mutations.

The validated gRNA for Cacna1g^MBH^ ^KO^ was cloned to AAV-U6-gRNA-CMV-SaCas9 vector by Vector Biolabs. The two validated gRNAs along with their U6 promoters for Cacna1g^POMC^ ^KO^ were cloned to pAAV-Linker-hSyn::mCherry.3xFLAG-WPRE (ref. Addgene #120391) in tandem with MluI and SacI sites respectively. The resultant plasmids along the gRNA-less empty vectors (as controls) were then provided to Vector Biolabs for AAV production.

The hCav3.1 from hα1Ga-pDsRed plasmid (ref; Addgene Cat# 45811) were subcloned to following vectors: 1.) pCAG-ires-GFP vector (Addgene #45025) with EcoRI and NotI sites for cell electrophysiology experiments; 2.) piRES-puro vector (Addgene Cat# 25728) with EcoRI and NotI sites for site-directed mutagenesis experiments.

The original hα1Ga-pDsRed plasmid was modified to include an HA epitope tag and a stop codon to the C-termianl using NotI and XbaI sites for the domain mapping experiment. The full length hCav3.1 were then re-cloned to this new vector without the DsRed sequence and in-frame with the HA tag using the EcoRI and NotI sites. The truncated hCav3.1 fragments (Domain I fragment: 1 – 735 aa; Domain II fragment: 736 – 1267 aa; Domain III fragment: 1268 – 1586 aa; Domain IV fragment: 1587 – 2314 aa) were cloned to this vector using FseI and NotI sites retaining the 32 N-terminal amino acid residues of full-length hCav3.1 which contains the secretion signal peptide and is also recognised by the anti-Cav3.1 antibody for immunoprecipitation in the fluorescent leucine binding assay.

Leucine binding hCav3.1 mutants were generated by performing site-directed mutagenesis using standard overlapping PCR method on the hα1Ga-pDsRed plasmid. The resultant hCav3.1 mutants were cloned into the piRES-puro vector using EcoRI and NotI sites. The sequence integrity of all plasmids was confirmed by Sanger sequencing (Eurofins Genomics).

### Indel mutation detection assay on genomic DNA

Genomic DNA was extracted from MBH tissues using TRIzol as described above and from cultured cells with QIAamp DNA kit (Qiagen). The T7 endonuclease I based mutation based EnGen Mutation Detection Kit (New England Biolabs) was used to detect on-target CRISPR-mediate indel mutations according to the manufacturer’s protocol (Version 3.0_7/20). The top exonic off-target loci were predicted using the CCTop online tool ^57^.

### Live cell fluo8 calcium flux assay

2.5×10^4^ cells per well were plated to poly-D-lysine coated 96 well plates with black wall and clear-bottom for 72hr and allowed to grow to confluence. After that, cells were rinsed once with imaging medium (CaCl_2_ 1.8 mM, MgCl_2_ 0.8 mM, KCl 5.4 mM, NaCl_2_ 115 mM, glucose 13.8 mM) and loaded with a cell permeable fluorescent calcium sensitive dye Fluo8-AM (Abcam) at 5 μM and allowed for 30 min incubation at 37°C. Imaging medium was then replaced (100 μl per well) and allowed for incubation at room temperature (∼24°C) in darkness for 30 min. Cells were then treated with indicated substances in a 5 μl volume. The plate was then sealed with a MicroAmp optical adhesive film (Thermo Fisher Scientific) and put into the Tecan infinite M1000Pro plate reader. After a short 5 second orbital shaking, fluorescence kinetics were measured under the bottle-reading mode with following settings: excitation: 490±9 nm, emission 520±9nm, number of Flashes: 10, flash frequency: 400 Hz, integration time: 20 us, interval time: 5 s, kinetic cycle: 150.

### In vitro fluorescent leucine and valine binding assay

An immunoprecipitation-based fluorescent leucine/valine binding assay was adapted from a radioactive labelled version ^20^. The HEK293-hCav3.1 stable cells were plated to a 6-well cell culture plate at 1 x 10^6^ per well density and allowed to grow to confluence for 48 hours. For the domain mapping experiment, HEK293 cells were transiently transfected with full-length and truncated hCav3.1 and allowed for expression for 48 hr as mentioned above. On the day of performing the binding assay, the antibody was conjugated to the magnetic protein A Dynabeads before cell harvesting. In brief, 33.3 μl beads per sample were washed with 1 mL TBS with 1% Triton X-100 (TBST) then blocked with 500 μl TBST with 1μg/ml BSA for 20 min at 4°C with rotation, followed by incubating with 1 μg N-terminal anti-Cav3.1 (Alomone Labs, Cat# ACC-021) (or 1 μg rabbit IgG as a control) in 500 μl lysis buffer with 1 μg/ml BSA at 4°C with rotation for 1 hr. After that, beads were washed twice with TBST.

Cells were harvested using a lysis buffer (TBS with 1% Triton X-100, 1x Roche EDTA free cOmplete protease inhibitor, 1x Roche pSTOP phosphatase inhibitors, 50mM sodium fluoride, 1mM sodium orthovanadate) and allowed to mix with gentle rotation at 4°C for 1 hr. After that, cells were centrifuged with 16100 rcf at 4°C for 10 min. Supernatant was then loaded to washed beads and allowed to incubate at 4°C with gentle rotation for 2 hr. Heat-denatured samples were obtained by heating the supernatant to 70°C for 10 min followed by chilling on ice for 5 min prior loading to beads. Then, the beads were washed 1 time with lysis buffer followed by 3 times high-salt TBST adjusted to 500 mM sodium chloride.

After the washing, beads were resuspended with 500 μl assay buffer (TBS with 0.1% Triton X-100) with 10 μM FAM-leucine or FAM-valine (Cambridge Research Biochemicals) and allowed to incubated at 4°C with gentle rotation for 1 hr. 10 mM unlabelled leucine/ valine were added to respective control samples at this stage to validate the specificity of the binding assay. The beads were then washed 3 times with 1 mL assay buffer. The immunocomplex was eluted from beads with 135 μl elution buffer (PBS with 1% SDS) and heating at 70°C for 10 min. The supernatant was then retrieved and 3 technical replicates per condition with 40 μl each were loaded to a 96 well plates with black wall and clear-bottom for fluorescence measurement the Tecan infinite M1000Pro plate reader (bottle-reading mode, excitation: 490±9 nm, emission 520±9 nm, number of Flashes: 10, flash frequency: 400 Hz, integration time: 20 μs). After the fluorescence measurement, the samples were retrieved and added 1X SDS sample-loading buffer for Western Blot analysis as described below.

### Co-immunoprecipitation

HEK293-hCav3.1 cells were transiently transfected with hTRPC5 plasmid (Sino Biological; Cat# HG18741-UT) using TransIT-293 Transfection Reagent. After 48 hr, cells harvested with an IP buffer (1% Triton X-100, 10% glycerol, 1x Roche cOmplete protease inhibitor cocktail with EDTA, in TBS pH7.4) at 4°C for 1 hr with gentle rotation and cleared by centrifugation. The cell lysates were subjected to anti-hCav3.1 IP by incubating with 1 μg of anti-hCav3.1 Rabbit antibody (Alomone) (or with 1 μg of unimmunised rabbit IgG (Thermo Fisher) as control) together with 30 μl of protein A agarose (Thermo Fisher) for 2 hr at 4°C with rotation. The immuno-complexes were washed 5 times with IP buffer and eluted by 1x SDS sample buffer and subjected to Western blot analysis as described above.

### In silico prediction of L-leucine binding sites on Cav3.1

To predict the possible binding site of L-Leucine onto Cav3.1, a blind docking approach using AutoDock Vina 1.1.2 was used as described previously ^58^. The cryo-EM derived structure of human Cav3.1 protein (PDB id: 6KZO) ^21^ as well as the 3D structures of the ligands (L-Leucine and L-Valine) were obtained from the Protein Data Bank. Prior to docking, the protein structure was prepared using the Dock Prep module of UCSF Chimera version 1.17.3 (https://www.cgl.ucsf.edu/chimera/). Initially, we performed blind docking of L-Leucine (the active ligand) and with L-Valine as a decoy against the entire Cav3.1 structure using AutoDock Vina at an exhaustiveness of 24 in three independent runs. Afterwards, total 27 docked poses (3 x 9/docking run) for each ligand were saved and clustered which allowed segregation into distinct potential sites or pockets. Through visual inspection, we then disregarded the sites where both Leucine and Valine poses overlapped. The sites that appeared to be potentially unique for L-Leucine recognition were then ranked based on the average of predicted energy of interaction (DG, kcol/mol) of the docked L-Leucine poses clustered per site. Finally, a focused docking of L-Leu was performed for each site using GOLD suite version 5.3 (CCDC, Cambridge) done in 15 independent runs/site. The most reproducible pose per site was then considered as the final, plausible binding mode of L-Leucine at each site. These poses in complex with the Cav3.1 protein were then subjected to 2D ligand interaction analysis through PLIP server ^59^ and depicted using MOE version 2019 (Chemical Computing Group, Montreal, Canada).

### Quantification and Statistical analysis

Unless otherwise stated, all the data were presented as means ± SEM and were analysed using GraphPad Prism 10 software package. For all the statistical tests, an α risk of 5% was used to define statistical significance. For pairwise comparisons were conducted using two-tailed t tests assuming equal variance between groups. One-factor multiple comparisons (across all groups or against control group) were tested with one-way ANOVAs followed by Holm-Sidak post hoc tests. For repeated measurement of the same subjects, paired t test or repeated measures one-way ANOVA were used. Two-factors multiple comparisons were tested with two-way ANOVAs followed by Holm-Sidak post hoc tests. All kinetics were analysed using repeated-measures two-way ANOVAs and adjusted with Holm-Sidak post hoc tests. Dietary and aCSF/leucine treatments were allocated randomly in weight-matched groups. We used blinding (to mouse genotype, viral treatment, or drug delivered) for in vivo experiments and to conduct image analyses. Additional statistical details on each experiment can be found in the figures or figure legends.

---

**Figure S1:**
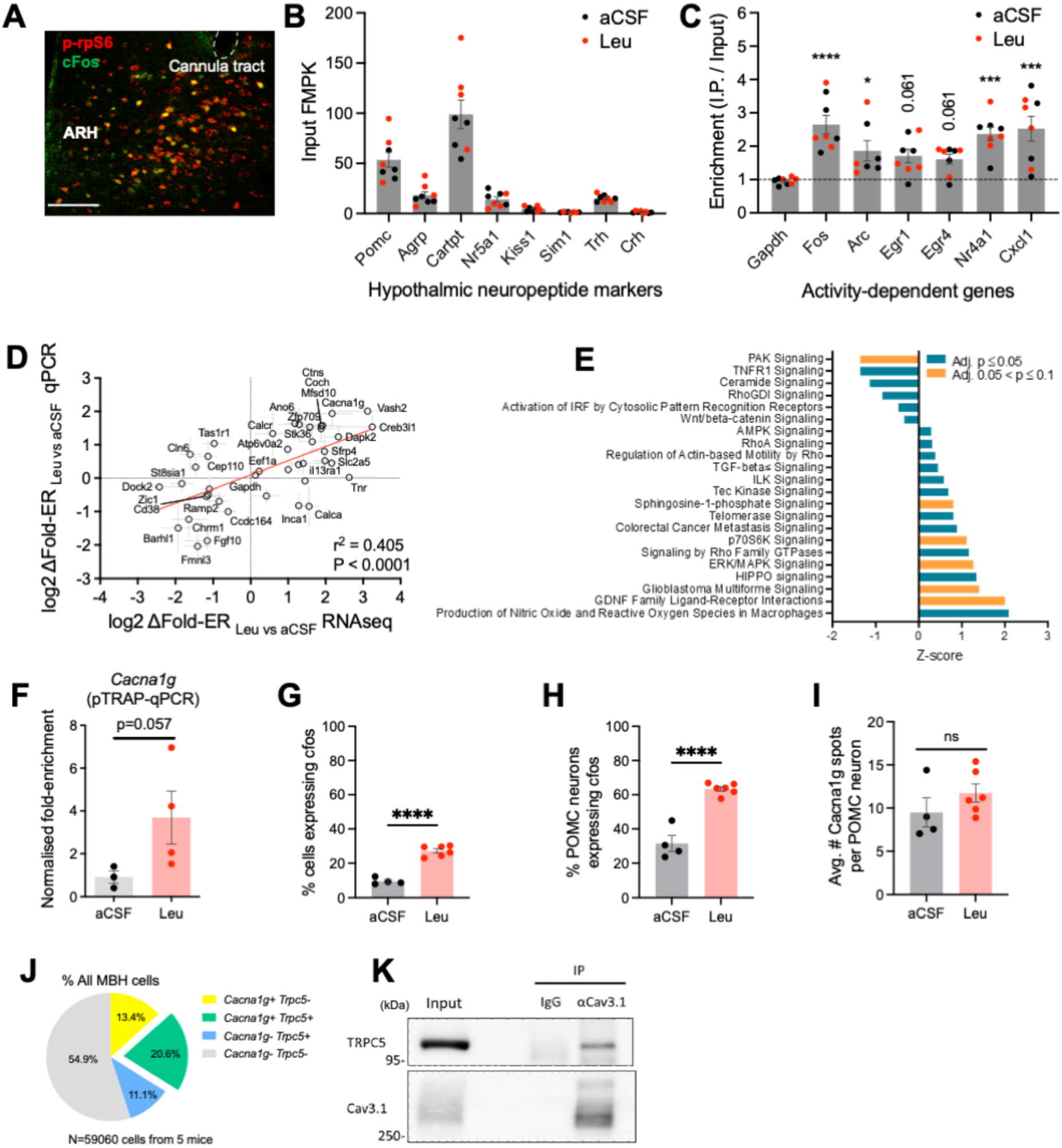
Additional data related to Figure 1. **(A)** Representative immunofluorescence confocal image showing extensive co-localisation of phosphorylated rpS6 and cFos in the ARH and VMH after MBH leucine injection. Scale bar: 50 μm. **(B)** Abundance of hypothalamic marker transcripts in MBH input samples from both aCSF and leucine groups. **(C)** Enrichment (I.P./Input) of activity dependent genes in MBH samples from both aCSF and leucine groups compared to *Gapdh*. **(D)** Validation of PhosphoTRAP RNAseq results with RT-qPCR of 41 selected genes. A statistically significant positive linear relationship is found between these two RNA quantification techniques. n = 4 for RNAseq, n = 3-4 for RT-qPCR. **(E)** Ingenuity Pathway analysis (IPA) of differentially enriched canonical pathways in leucine-responsive neurons identified in PhosphoTRAP assay. See Table S5 for full data. **(F)** Normalised fold enrichment ratio of *Cacna1g* analysed by RT-qPCR. The RNA samples used were from the same experiment also submitted for RNAseq. n = 3 for aCSF; n = 4 for Leu. **(G-I)** Quantification RNAscope analysis after MBH leucine injection as shown in (Figure 1C). (**G**) % all MBH cells expressing *cfos*, (**H**) % all POMC neurons expressing *cfos*, (**I**) number of *Cacna1g* spots per POMC neuron. n = 4 for aCSF; n = 6 for Leu. **(J)** Relative proportion of all MBH cells expressing *Cacna1g* and *Trpc5*. N = 59060 MBH cells from 5 WT mice. **(K)** Representative Western blot of co-immunoprecipitation assay of heterologously expressed human TRPC5 and Cav3.1 in HEK293 cells. This experiment has been repeated twice with similar results. *p<0.05, ***p<0.001, ****p<0.0001. Values are reported as mean ± SEM

**Figure S2:**
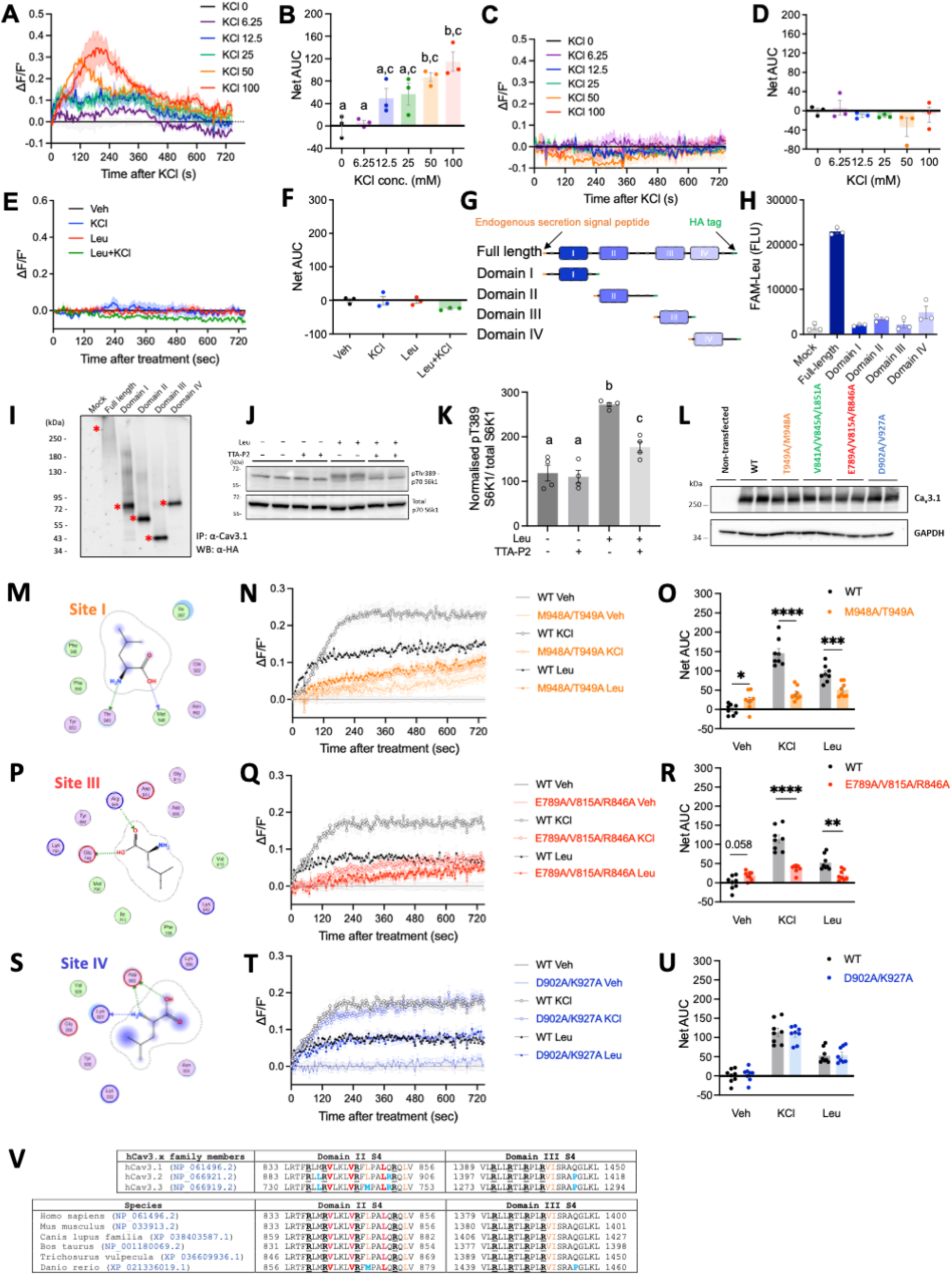
Additional data related to Figure 3 **(A-F)** Fluo8 calcium flux assay of HEK293 (-hCa_v_3.1 and -WT) cells. Dose-dependent KCl-induced calcium responses and AUC quantification of HEK293-hCa_v_3.1 (**A-B**); Lack of calcium responses in non-transfected HEK293-WT cells after various doses of KCl (**C-D**) and leucine treatment (**E-F**). n = 3 per group. **(G-I)** In vitro fluorescent leucine binding assay of individual hCa_v_3.1 domains. Schematic diagram of recombinant truncated hCav3.1 domain fragments construction (**G**). Quantification of fluorescence measurements for fluorescent leucine binding (**H**). Post-assay Western blot validates successful expression and immunoprecipitation of the recombinant domain fragments (**I**). Red asterisks indicate expected band size of the fragments. Note that this assay was performed on transiently transfected cells; the full-length hCa_v_3.1 expression was far weaker yet bound stronger to FAM-leucine compared to the truncated fragments. Data represent 3 technical replicates upon fluorescence measurements. These experiments have been repeated twice with similar results. Data of a representative experiment is shown. **(J-K)** Cav3.1 contributes to leucine induced mTOR1 activation in HEK293-hCav3.1 cells. (**J**) Representative Western blot of Thr389 phosphorylation of S6K in HEK293-hCav3.1 cells treated with leucine. with or without co-treatment of TTA-P2. (**K**) Normalised densitometric quantification of pThr389 S6K to total S6K ratio from the same blot. Data were merged from 2 independent experiments with duplicates each. **(L)** Western blot validation for the protein expression of hCav3.1 Site I/II/III/IV mutants in respective stably expressing HEK293 cells. **(M, P, S)** 2D ligand interactions diagrams for the docked poses of L-Leu at Site I (**M**), Site III (**P**) and Site IV (**S**). The residues mutated for experimental validation are shown with asterisks. The dotted arrow signs indicate hydrogen bonding whilst the dotted contour around the ligand pose indicate hydrophobic interactions with residues shown. **(N-O, Q-R, T-U)** Fluo8 calcium flux assay of HEK293-hCa_v_3.1 predicted Leu-binding mutant cells. Calcium responses and AUC quantification after KCl and Leu treatments of Site I mutant cells (M948A/T949A; **N-O**); Site III mutant cells (E789A/V815A/R846A; **Q-R**) and Site IV mutant cells (D902A/K927A; **T-U**). n = 8 merged from 4 independent experiments with duplicates each. **(V)** Amino acid sequence alignments of the domain II and III voltage-sensing S4 transmembrane segments of human Cav3.1 against hCav3.2 and hCav3.3 (upper panel) and against Cav3.1 of various mammalian species and zebrafish (lower panel). Bolded underlined black “R”: consensus voltage-sensing arginine residues; bolded red: critical residues predicted to mediate leucine binding and were chosen for mutagenesis in this study; orange: other residues predicted to participate in leucine binding; bolded blue: residues different from human Cav3.1. Groups denoted with different letter in (B,L) indicate significant difference (p<0.05). *p<0.05, **p<0.01, ***p<0.01, ****p<0.0001. Values and calcium response traces in (A, C, E, N, Q, T) are reported as mean ± SEM.

**Figure S3:**
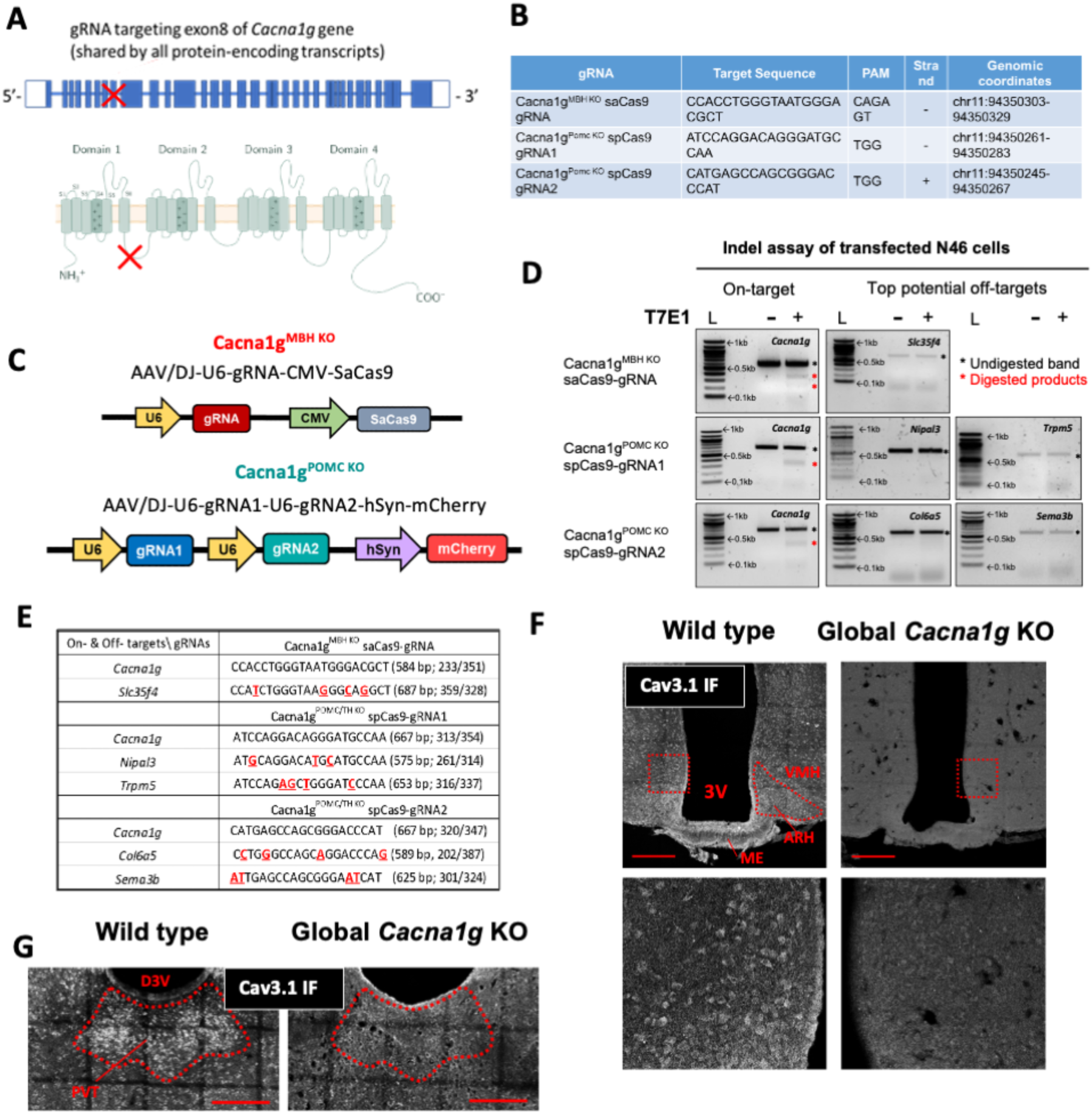
CRISPR-mediated *Cacna1g* KO strategy, related to Figures 4-5 **(A)** Diagram of the CRISPR knockout strategy targeting the exon 8 of mouse *Cacna1g* gene which is common to all protein-coding splice variants. The red crosses indicate the intended mutation sites on genomic DNA and protein. **(B)** Detail information of the gRNAs used in this study. **(C)** Schematic representation of the AAV constructs used for Cacna1g^MBH^ ^KO^ and Cacna1g^POMC^ ^KO^ models. **(D)** Validation for the specificity of gRNAs used to target mouse *Cacna1g.* Representative DNA electrophoresis images of T7 endonuclease I mutation detection assay of genomic DNA isolated from mouse mHypoE-N46 cell line 96 hr after transfection of indicated gRNA/Cas9 all-in-one plasmids. Intended *Cacna1g* targeting site and top exonic off-target sites predicted by CCTop web software were submitted to the analysis. Black arrows indicate parental unmodified PCR amplicons and red arrows indicate digested mutated fragments of expected size. L: 0.5 μg 100 bp DNA ladder; +/-: with or without T7E1 digestion. Note that images were adjusted differently to facilitate visualisation of fainter bands. **(E)** Table summary of the on- and predicted off-targets genomic DNA sequence for the gRNAs used in this study. The expected DNA size of parental amplicons and digested fragments are indicated in the brackets. **(F-G)** Confocal immunofluorescence (IF) microscopy validation of a C-terminal specific anti-Cav3.1 antibody (Abcam, Cat# ab203577) with constitutive global *Cacna1g* KO mice. (**F**) Representative IF images of the MBH of WT and global KO mice. Upper panel: multi-focal plane (maximum intensity projection) images. Lower panel: single focal plane zoom-in images of the regions highlighted in red squares of the respective pictures shown on the upper panel. (**G**) Representative IF images of the thalamus of WT and global *Cacna1g* KO mice. 3V: third ventricle; ARH: arcuate nucleus of the hypothalamus; VMH: ventromedial nucleus of the hypothalamus; ME: median eminence; D3V: dorsal third ventricle; PVT: paraventricular nucleus of the thalamus. Scale bars: 200 μm.

**Figure S4:**
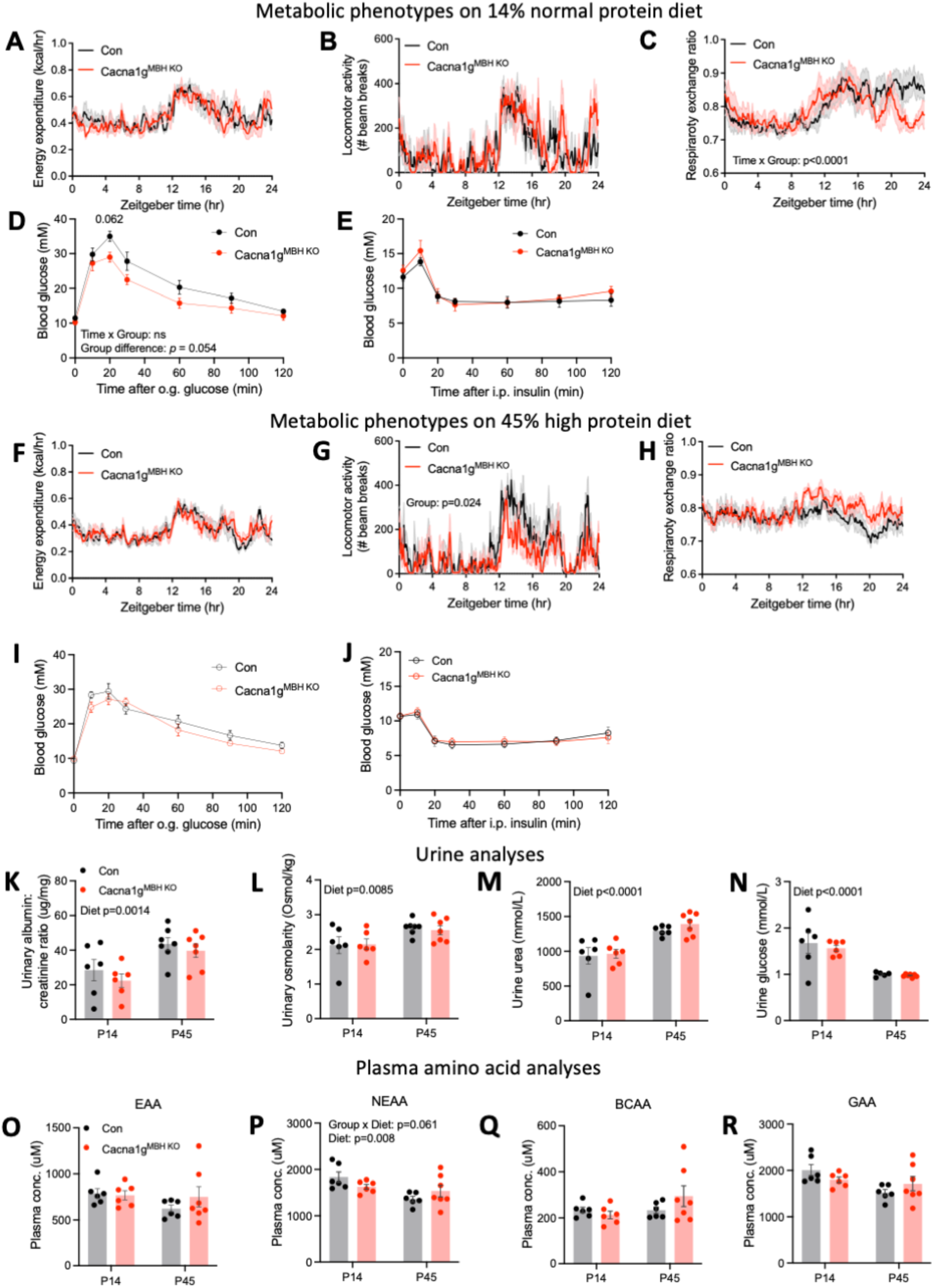
Metabolic and physiologic parameters of Cacna1g^MBH^ ^KO^ mice, related to Figure 4 **(A-C)** Metabolic parameters of Con and Cacna1g^MBH^ ^KO^ mice on P14 diet during indirect calorimetry measurement. (**A**) 24 hr profiles of energy expenditure, (**B**) respiratory exchange ratio and (**C**) locomotor activity. **(D-E)** Glucose homeostasis regulation of Con and Cacna1g^MBH^ ^KO^ mice on P14 diet. (**D**) Oral glucose tolerance test and (**E**) insulin tolerance test. **(F-H)** Metabolic parameters of Con and Cacna1g^MBH^ ^KO^ mice on P45 diet during indirect calorimetry measurement. (**F**) 24 hr profiles of energy expenditure, (**G**) respiratory exchange ratio and (**H**) locomotor activity. **(I-J)** Glucose homeostasis regulation of Con and Cacna1g^MBH^ ^KO^ mice on P45 diet. (**I**) Oral glucose tolerance test and (**J**) insulin tolerance test. **(K-N)** Urine analyses of Con and Cacna1g^MBH^ ^KO^ mice fed on P14 and P45 diets. Urinary albumin/creatinine (ACR) ratio (**K**), osmolarity (**L**), urea (**M**) and glucose (**N**). **(O – R)** Grouped post-absorptive phase plasma amino acid profiling of Con and Cacna1g^MBH^ ^KO^ mice fed on P14 and P45 diets. Essential amino acid (EAA; **O**), non-essential amino acid excluding glycine (NEAA; **P**), glucogenic amino acid excluding glycine (GAA, **Q**), branched-chain amino acid (BCAA, **R**). Individual amino acid data are shown in Table S7.Con-P14: n = 6; Con-P45: n = 6; KO-P14: n = 7; KO-P45: n = 7. Values are reported as mean ± SEM.

**Figure S5:**
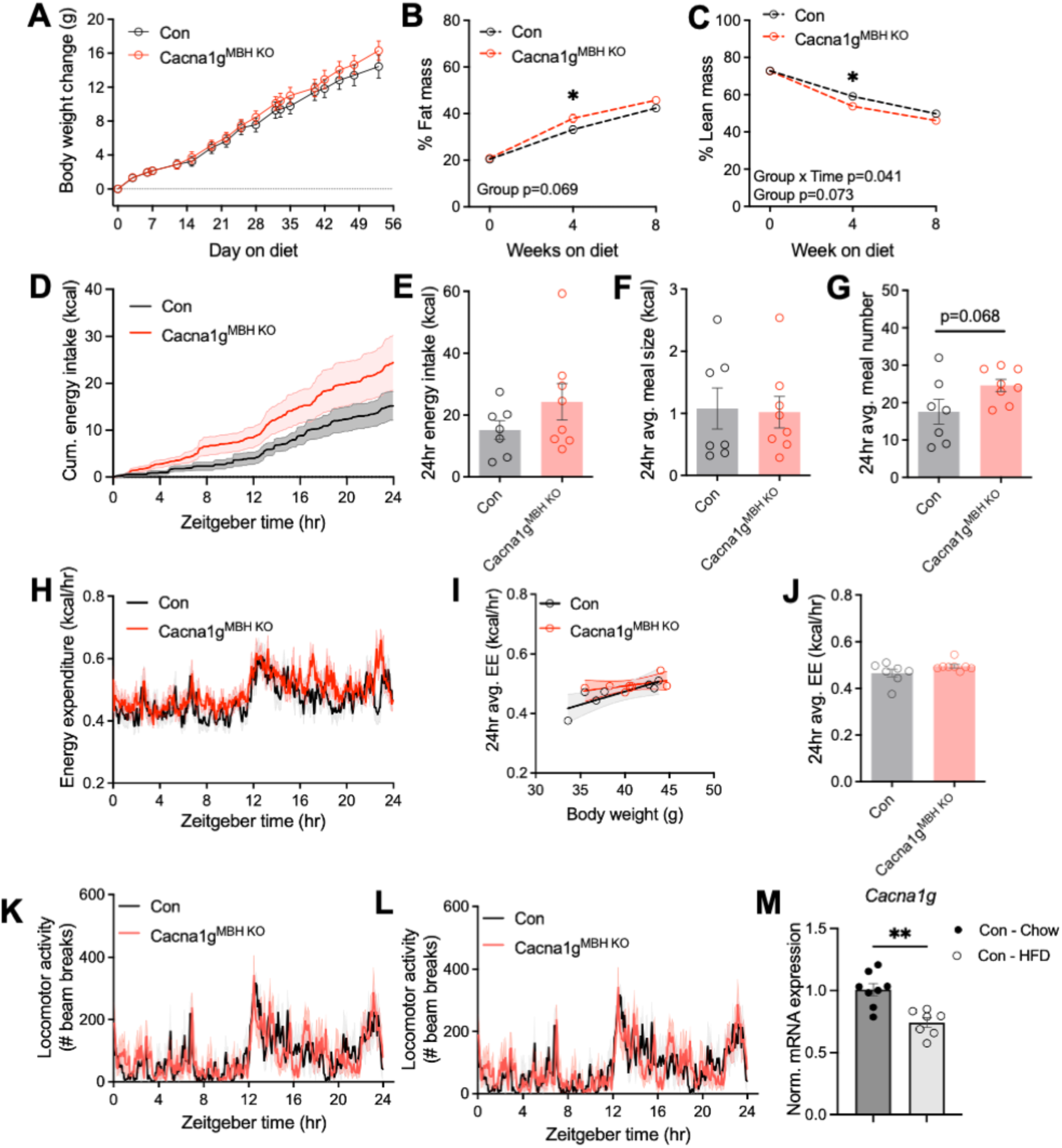
Hypothalamic *Canca1g* is downregulated by diet-induced obesity, related Figure 4 **(A-L)** Metabolic phenotyping of Con and Cacna1g^MBH^ ^KO^ mice on 60% high fat diet (HFD). (**A**) Body weight change, (**B**) % fat mass and (**C**) % lean mass over the course of 8 weeks HFD feeding. (**D**) 24 hr energy intake profile, (**E**) 24 hr total energy intake, (**F**) 24 hr average meal size, (**G**) 24 hr average meal number, (**H**) 24 hr profiles of energy expenditure, (**I**) ANCOVA analysis of 24 hr average energy expenditure against body weight, (**J**) 24 hr average energy expenditure, (**K**) 24 hr profile of locomotor activity, (**L**) 24 hr profile of respiratory exchange ratio, during indirect calorimetry measurement. **(M)** RT-qPCR measurement of relative expression of *Cacna1g* mRNA in the mediobasal hypothalamus of Con mice fed on HFD vs chow for 8 weeks.Con-Chow: n = 8; Con-HFD: n = 7; KO-Chow: n = 7; KO-HFD: n = 8. *p<0.05. **p<0.01. Values are reported as mean ± SEM.

**Figure S6:**
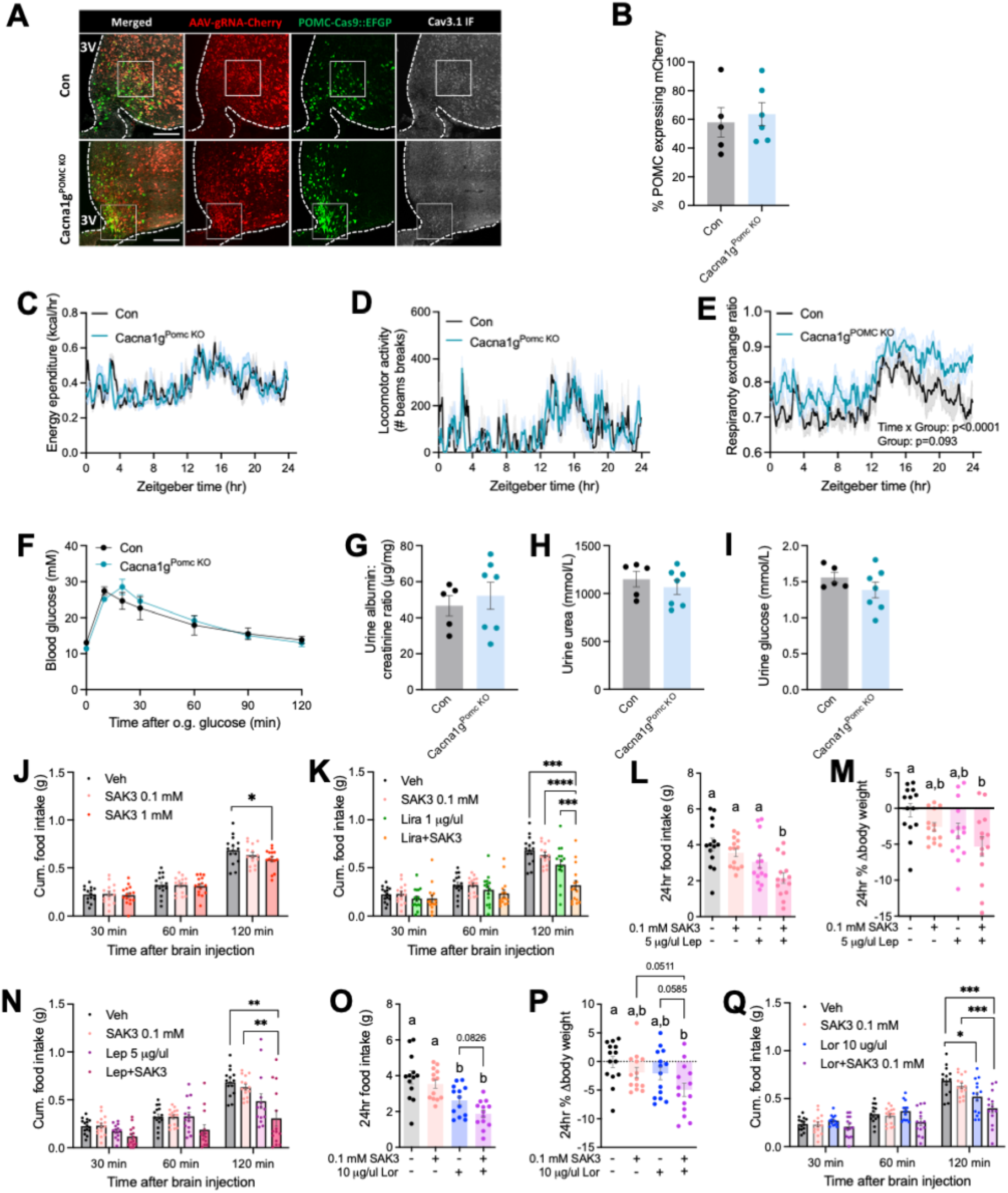
Additional data related to Figure 5-6 **(A-B)** Confocal **i**mmunofluorescence (IF) microscopy analysis of the CRISPR knockout efficiency of Cav3.1 in Cacna1g^POMC^ ^KO^ mediobasal hypothalamus with a C-terminal specific Cav3.1 antibody validated in Fig. S3F. (**A**) Zoom-out images of Fig. 5B. Scale bars: 200 µm. (**B**) Quantification of hypothalamic POMC neurons transduced with AAVs. Con: n = 5; Cacna1g^POMC^ ^KO^: n = 6. **(C-E)** Metabolic parameters of Cacna1g^POMC^ ^KO^ mice on P45 diet. (**C**) 24 hr profile of energy expenditure, (**C**) 24 hr profiles of locomotor activity and (**D**) respiratory exchange ratio (**E**) during indirect calorimetry measurement. Con: n = 5; Cacna1g^POMC^ ^KO^: n = 7. **(F)** Oral glucose tolerance test of Con and Cacna1g^MBH KO^ mice on P14 diet. Con: n = 5; Cacna1g^POMC^ ^KO^: n = 7. **(G-I)** Urine analyses of Con and Cacna1g^POMC^ ^KO^ mice fed on P45 diets. Urinary albumin/creatinine ACR ratio (**G**), urea (**H**) and glucose (**I**). Con: n = 5; Cacna1g^POMC^ ^KO^: n = 7. **(J)** Acute feeding responses after MBH injection of Veh, 0.1 mM and 1 mM Ca_v_3.1 activator SAK3. n = 14 per group. **(K)** Acute feeding responses after MBH injection of Veh, 0.1 mM SAK3, 1 µg/μl liraglutide (Lira) and SAK3+Lira co-treatment. n = 14 per group. **(L-N)** 24 hr feeding responses (**L**), 24 hr % body weight changes (**M**) and acute feeding responses (**N**) after MBH injection of Veh, 0.1 mM SAK3, 5ug/μl leptin (Lep) and SAK3+Lep co-treatment. n = 14 per group. **(O-Q)** 24 hr feeding responses (**O**), 24 hr % body weight changes (**P**) and acute feeding responses (**Q**) after MBH injection of Veh, 0.1 mM SAK3, 10ug/μl lorcaserin (Lor) and SAK3+Lor co-treatment. n = 14 per group. Groups denoted with different letter in (L, M, O, P) indicate significant difference (p<0.05). *p<0.05, **p<0.01, ***p<0.001, ****p<0.0001. Values are reported as mean ± SEM.

