## Supplementary figures for "Cav3.1 is a leucine sensor in POMC neurons mediating appetite suppression and weight loss"

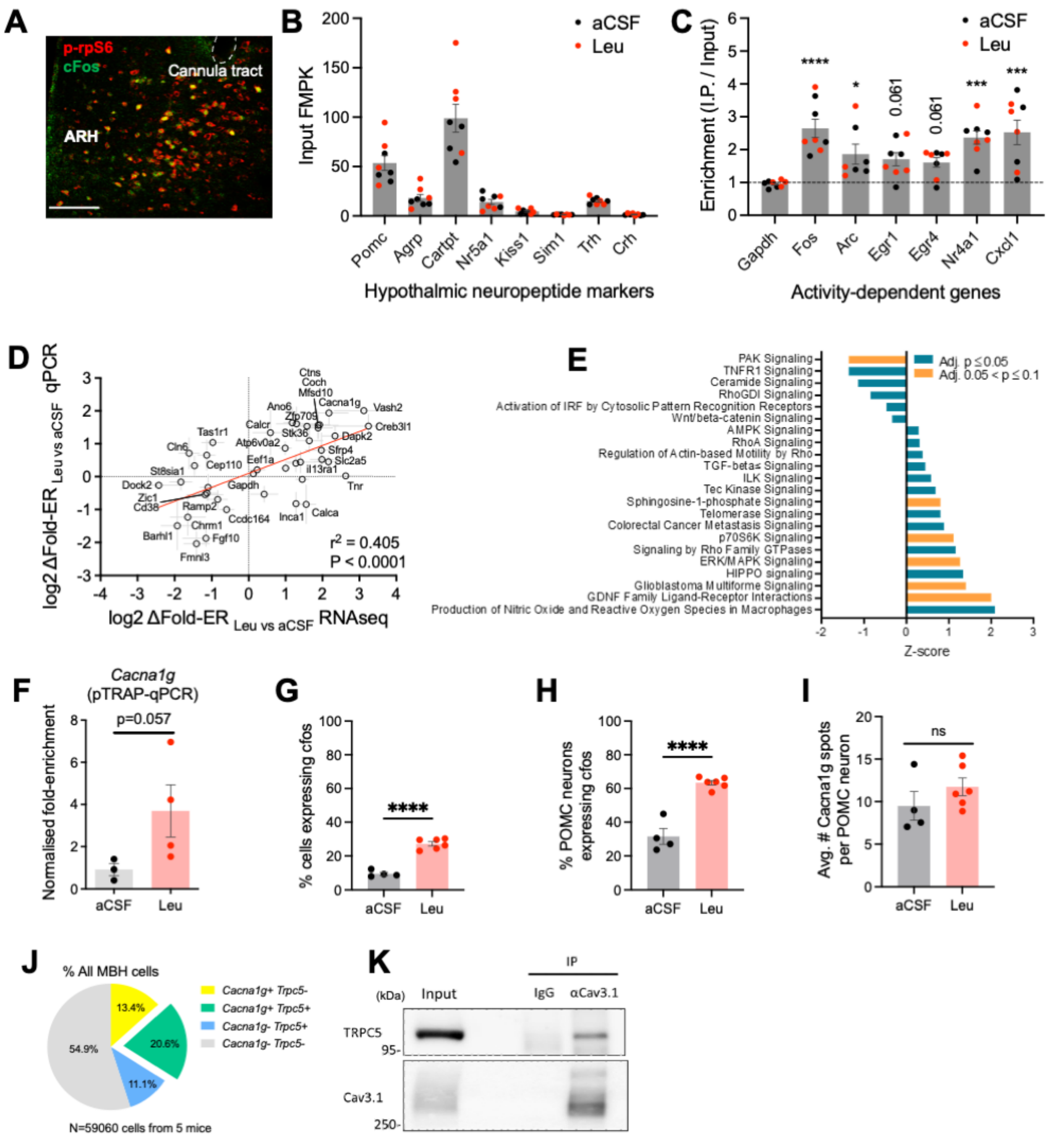


**Figure S1: Additional data related to Figure 1.**

**(A)** Representative immunofluorescence confocal image showing extensive co-localisation of phosphorylated rpS6 and cFos in the ARH and VMH after MBH leucine injection. Scale bar: 50 µm. **(B)** Abundance of hypothalamic marker transcripts in MBH input samples from both aCSF and leucine groups. **(C)** Enrichment (I.P./Input) of activity dependent genes in MBH samples from both aCSF and leucine groups compared to *Gapdh*. **(D)** Validation of PhosphoTRAP RNAseq results with RT-qPCR of 41 selected genes. A statistically significant positive linear relationship is found between these two RNA quantification techniques. n = 4 for RNAseq, n = 3-4 for RT-qPCR. **(E)** Ingenuity Pathway analysis (IPA) of differentially enriched canonical pathways in leucine-responsive neurons identified in PhosphoTRAP assay. See Table S5 for full data. **(F)** Normalised fold enrichment ratio of *Cacna1g* analysed by RT-qPCR. The RNA samples used were from the same experiment also submitted for RNAseq. n = 3 for aCSF; n = 4 for Leu. **(G-I)** Quantification RNAscope analysis after MBH leucine injection as shown in (Figure 1C). (**G**) % all MBH cells expressing *cfos*, (**H**) % all POMC neurons expressing *cfos*, (**I**) number of *Cacna1g* spots per POMC neuron. n = 4 for aCSF; n = 6 for Leu. **(J)** Relative proportion of all MBH cells expressing *Cacna1g* and *Trpc5*. N = 59060 MBH cells from 5 WT mice. **(K)**

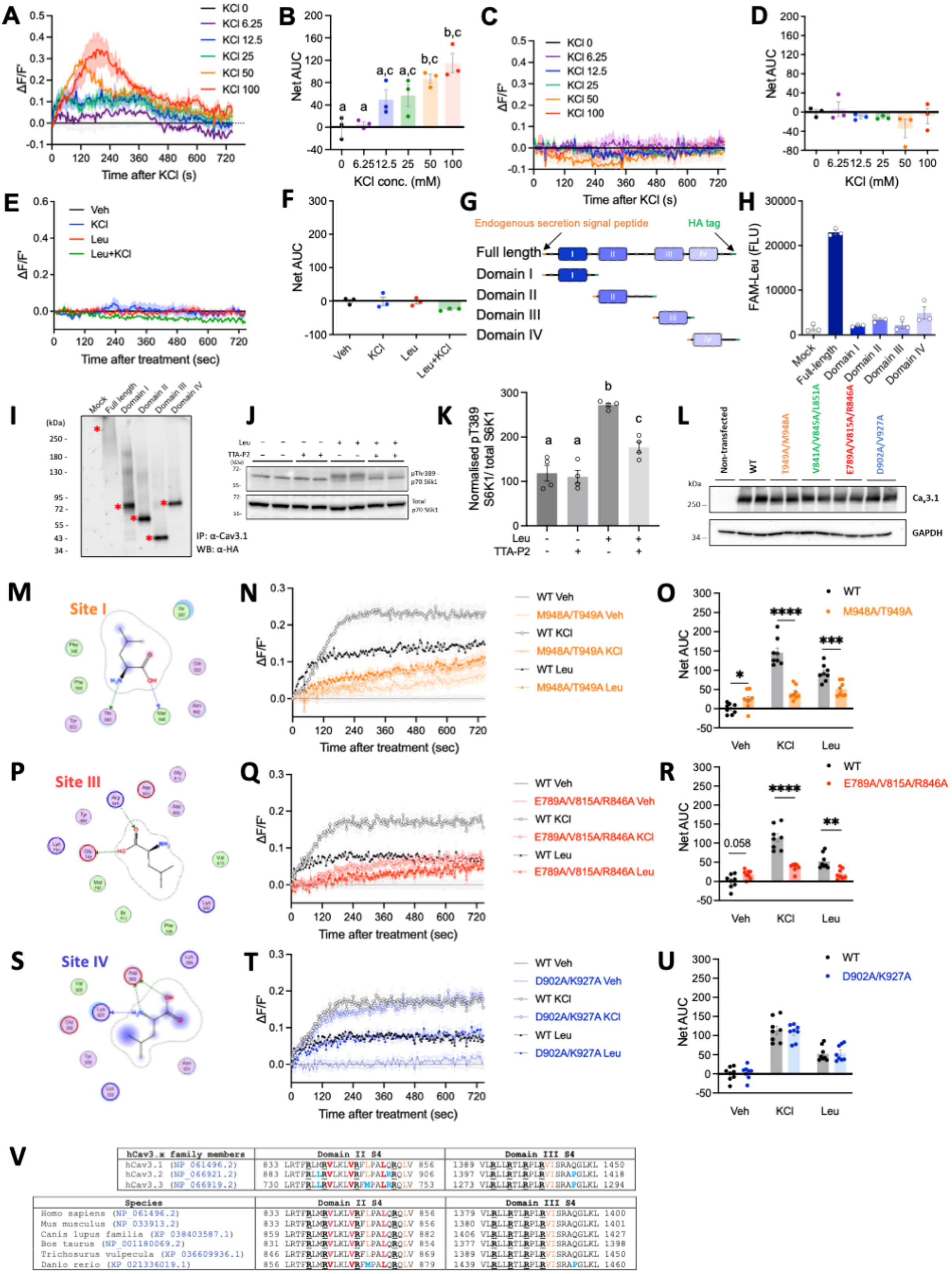


**Figure S2: Additional data related to Figure 3**

**(A-F)** Fluo8 calcium flux assay of HEK293 (-hCa$_v$3.1 and -WT) cells. Dose-dependent KCl-induced calcium responses and AUC quantification of HEK293-hCa$_v$3.1 (**A-B**); Lack of calcium responses in non-transfected HEK293-WT cells after various doses of KCl (**C-D**) and leucine treatment (**E-F**). n = 3 per group. **(G-I)** In vitro fluorescent leucine binding assay of individual hCa$_v$3.1 domains. Schematic diagram of recombinant truncated hCav3.1 domain fragments construction (**G**). Quantification of fluorescence measurements for fluorescent leucine binding (**H**). Post-assay Western blot validates successful expression and immunoprecipitation of the recombinant domain fragments (**I**). Red asterisks indicate expected band size of the fragments. Note that this assay was performed on transiently transfected cells; the full-length hCa$_v$3.1 expression was far weaker yet bound stronger to FAM-leucine compared to the truncated fragments. Data represent 3 technical replicates upon fluorescence measurements. These experiments have been repeated twice with similar results. Data of a representative experiment is shown. **(J-K)** Cav3.1 contributes to leucine induced mTOR1 activation in HEK293-hCav3.1 cells. (**J**) Representative Western blot of Thr389 phosphorylation of S6K in HEK293-hCav3.1 cells treated with leucine. with or without co-treatment of TTA-P2. (**K**) Normalised densitometric quantification of pThr389 S6K to total S6K ratio from the same blot. Data were merged from 2 independent experiments with duplicates each. **(L)** Western blot validation for the protein expression of hCav3.1 Site I/II/III/IV mutants in respective stably expressing HEK293 cells. **(M, P, S)** 2D ligand interactions diagrams for the docked poses of L-Leu at Site I (**M**), Site III (**P**) and Site IV (**S**). The residues mutated for experimental validation are shown with asterisks. The dotted arrow signs indicate hydrogen bonding whilst the dotted contour around the ligand pose indicate hydrophobic interactions with residues shown. **(N-O, Q-R, T-U)** Fluo8 calcium flux assay of HEK293-hCa$_v$3.1 predicted Leu-binding mutant cells. Calcium responses and AUC quantification after KCl and Leu treatments of Site I mutant cells (M948A/T949A; **N-O**); Site III mutant cells (E789A/V815A/R846A; **Q-R**) and Site IV mutant cells (D902A/K927A; **T-U**). n = 8 merged from 4 independent experiments with duplicates each. **(V)** Amino acid sequence alignments of the Domain II and III voltage-sensing S4 transmembrane segments of human Cav3.1 against hCav3.2 and hCav3.3 (upper panel) and against Cav3.1 of various mammalian species and zebrafish (lower panel). Bolded underlined black "R": consensus voltage-sensing arginine residues; bolded red: critical residues predicted to mediate leucine binding and were chosen for mutagenesis in this study; orange: other residues predicted to participate in leucine binding; bolded blue: residues different from human Cav3.1. Groups denoted with different letter in (B,L) indicate significant difference (p<0.05). *p<0.05, **p<0.01, ***p<0.01, ****p<0.0001. Values and calcium response traces in (A, C, E, N, Q, T) are reported as mean ± SEM.

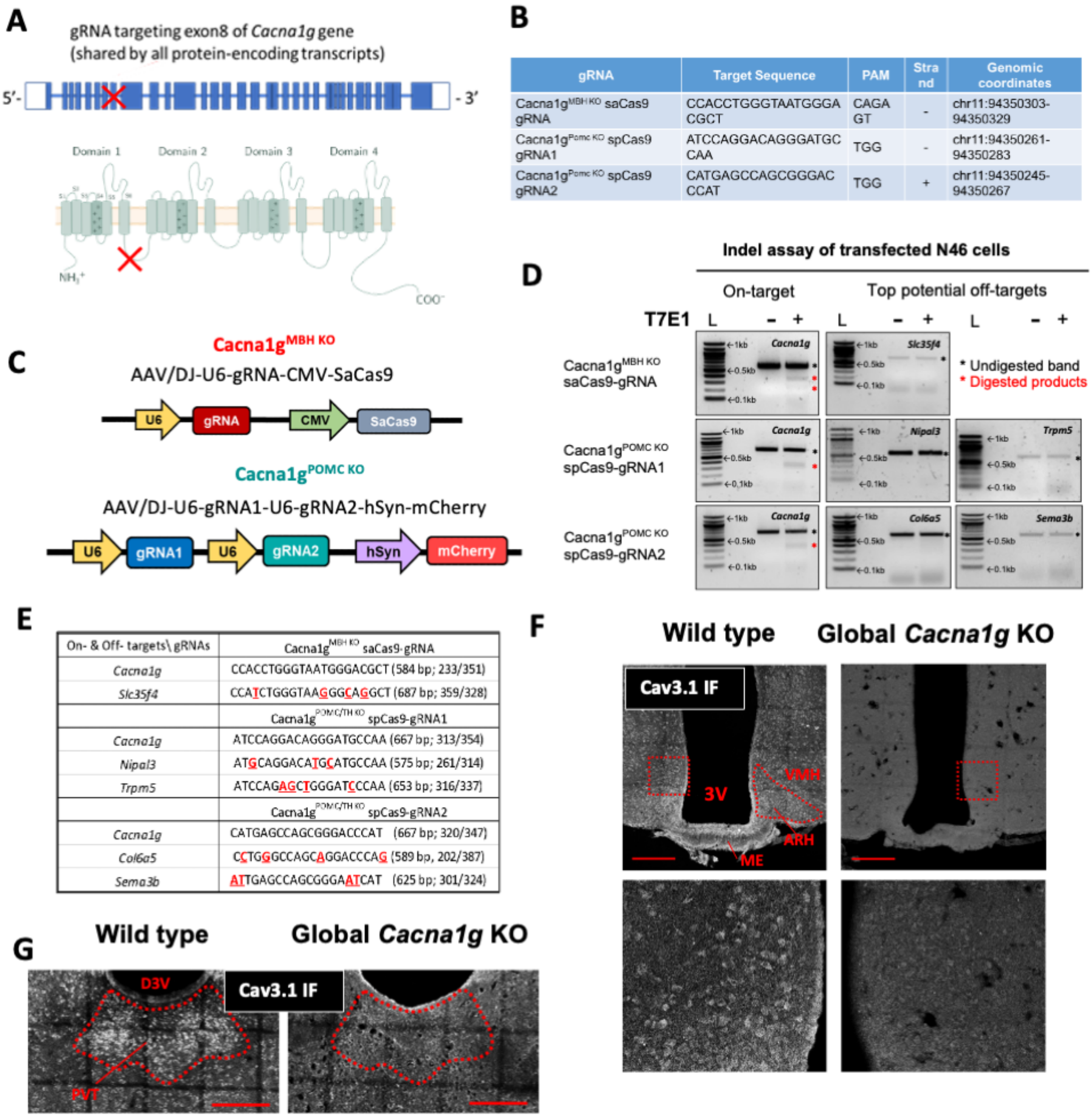


**Figure S3: CRISPR-mediated *Cacna1g* KO strategy, related to Figures 4-5**

**(A)** Diagram of the CRISPR knockout strategy targeting the exon 8 of mouse *Cacna1g* gene which is common to all protein-coding splice variants. The red crosses indicate the intended mutation sites on genomic DNA and protein. **(B)** Detail information of the gRNAs used in this study. **(C)** Schematic representation of the AAV constructs used for Cacna1g$^{MBH\ KO}$ and Cacna1g$^{POMC\ KO}$ models. **(D)** Validation for the specificity of gRNAs used to target mouse *Cacna1g.* Representative DNA electrophoresis images of T7 endonuclease I mutation detection assay of genomic DNA isolated from mouse mHypoE-N46 cell line 96 hr after transfection of indicated gRNA/Cas9 all-in-one plasmids. Intended *Cacna1g* targeting site and top exonic off-target sites predicted by CCTop web software were submitted to the analysis. Black arrows indicate parental unmodified PCR amplicons and red arrows indicate digested mutated fragments of expected size. L: 0.5 µg 100 bp DNA ladder; +/-: with or without T7E1 digestion. Note that images were adjusted differently to facilitate visualisation of fainter bands. **(E)** Table summary of the on- and predicted off-targets genomic DNA sequence for the gRNAs used in this study. The expected DNA size of parental amplicons and digested fragments are indicated in the brackets. **(F-G)** Confocal immunofluorescence (IF) microscopy validation of a C-terminal specific anti-Cav3.1 antibody (Abcam, Cat# ab203577) with constitutive global *Cacna1g* KO mice. (**F**) Representative IF images of the MBH of WT and global KO mice. Upper panel: multi-focal plane (maximum intensity projection) images. Lower panel: single focal plane zoom-in images of the regions highlighted in red squares of the respective pictures shown on the upper panel. (**G**) Representative IF images of the thalamus of WT and global *Cacna1g* KO mice. 3V: third ventricle; ARH: arcuate nucleus of the hypothalamus; VMH: ventromedial nucleus of the hypothalamus; ME: median eminence; D3V: dorsal third ventricle; PVT: paraventricular nucleus of the thalamus. Scale bars: 200 µm.

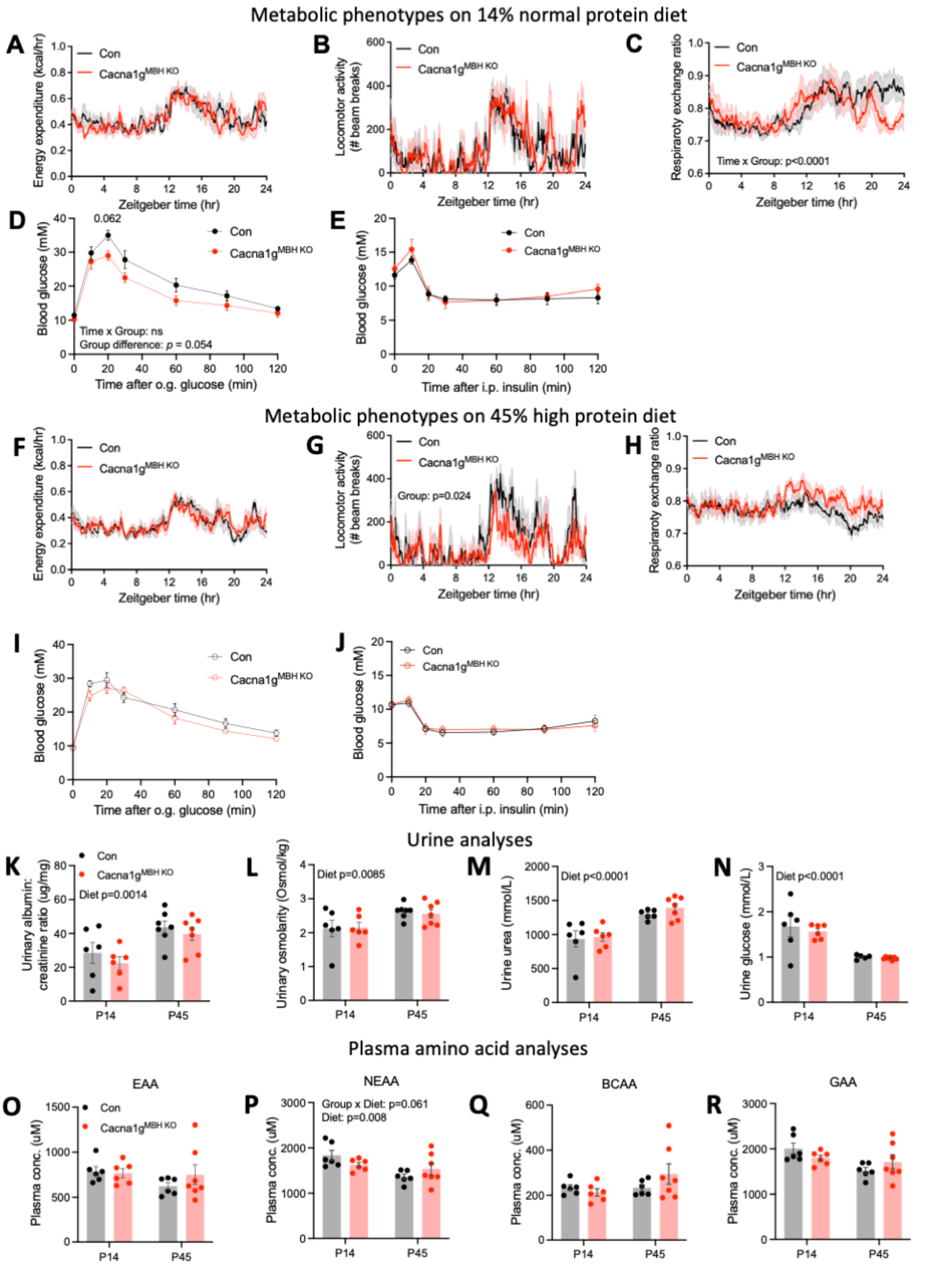


**Figure S4: Metabolic and physiologic parameters of Cacna1g$^{\mathrm{MBH\ KO}}$ mice, related to Figure 4**
**(A-C)** Metabolic parameters of Con and Cacna1g$^{\mathrm{MBH\ KO}}$ mice on P14 diet during indirect calorimetry measurement. (**A**) 24 hr profiles of energy expenditure, (**B**) respiratory exchange ratio and (**C**) locomotor activity. **(D-E)** Glucose homeostasis regulation of Con and Cacna1g$^{\mathrm{MBH\ KO}}$ mice on P14 diet. (**D**) Oral glucose tolerance test and (**E**) insulin tolerance test. **(F-H)** Metabolic parameters of Con and Cacna1g$^{\mathrm{MBH\ KO}}$ mice on P45 diet during indirect calorimetry measurement. (**F**) 24 hr profiles of energy

expenditure, (**G**) respiratory exchange ratio and (**H**) locomotor activity. **(I-J)** Glucose homeostasis regulation of Con and Cacna1g$^{MBH\ KO}$ mice on P45 diet. (**I**) Oral glucose tolerance test and (**J**) insulin tolerance test. **(K-N)** Urine analyses of Con and Cacna1g$^{MBH\ KO}$ mice fed on P14 and P45 diets. Urinary albumin/creatinine (ACR) ratio (**K**), osmolarity (**L**), urea (**M**) and glucose (**N**). **(O – R)** Grouped post-absorptive phase plasma amino acid profiling of Con and Cacna1g$^{MBH\ KO}$ mice fed on P14 and P45 diets. Essential amino acid (EAA; **O**), non-essential amino acid excluding glycine (NEAA; **P**), glucogenic amino acid excluding glycine (GAA, **Q**), branched-chain amino acid (BCAA, **R**). Individual amino acid data are shown in Table S7.Con-P14: n = 6; Con-P45: n = 6; KO-P14: n = 7; KO-P45: n = 7. Values are reported as mean ± SEM.

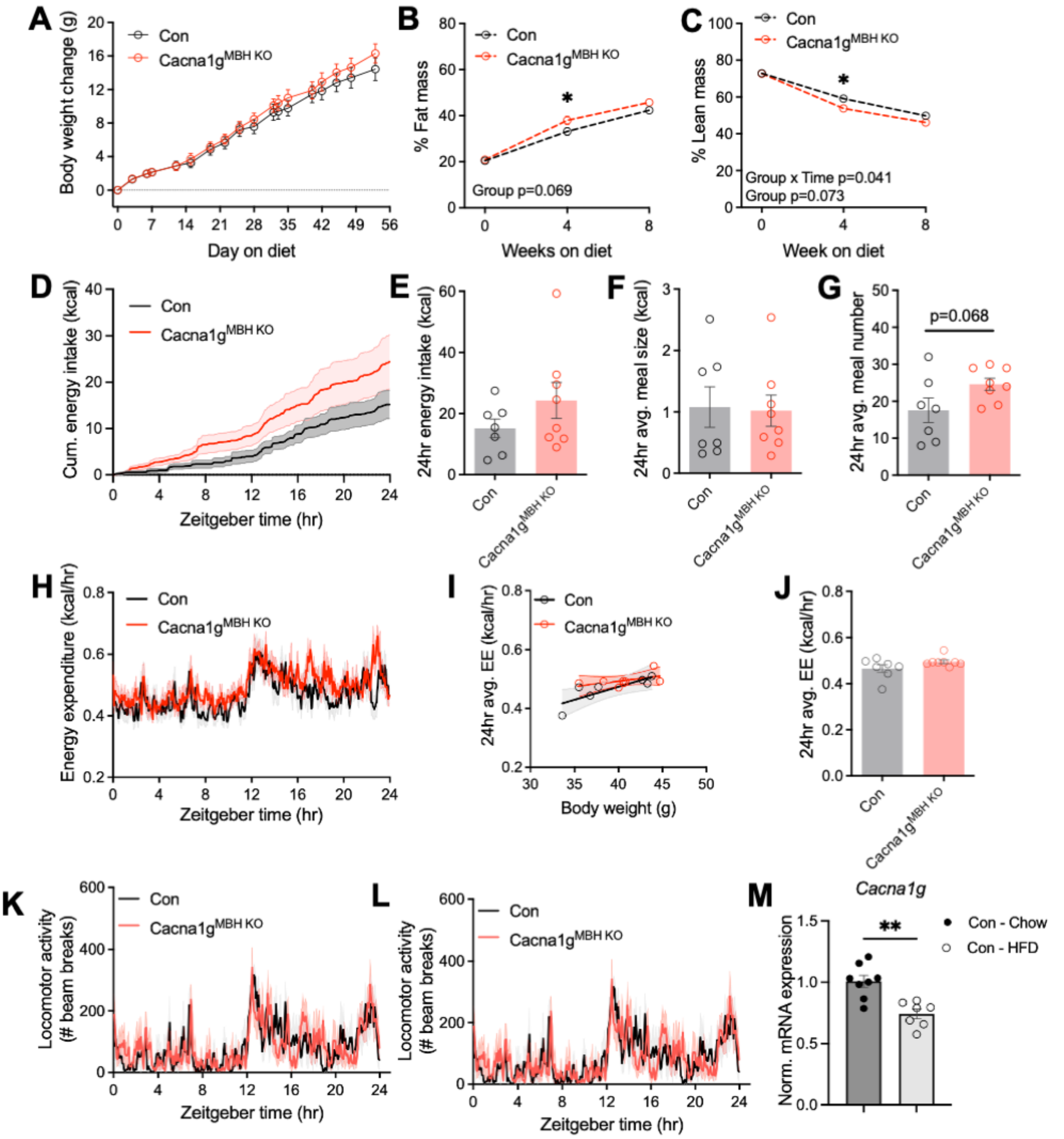


**Figure S5: Hypothalamic *Canca1g* is downregulated by diet-induced obesity, related Figure 4**
**(A-L)** Metabolic phenotyping of Con and Cacna1g$^{MBH\ KO}$ mice on 60% high fat diet (HFD). (**A**) Body weight change, (**B**) % fat mass and (**C**) % lean mass over the course of 8 weeks HFD feeding. (**D**) 24 hr energy intake profile, (**E**) 24 hr total energy intake, (**F**) 24 hr average meal size, (**G**) 24 hr average meal number, (**H**) 24 hr profiles of energy expenditure, (**I**) ANCOVA analysis of 24 hr average energy expenditure against body weight, (**J**) 24 hr average energy expenditure, (**K**) 24 hr profile of locomotor activity, (**L**) 24 hr profile of respiratory exchange ratio, during indirect calorimetry measurement. **(M)** RT-qPCR measurement of relative expression of *Cacna1g* mRNA in the mediobasal hypothalamus of Con mice fed on HFD vs chow for 8 weeks.Con-Chow: n = 8; Con-HFD: n = 7; KO-Chow: n = 7; KO-HFD: n = 8. *p<0.05. **p<0.01. Values are reported as mean ± SEM.

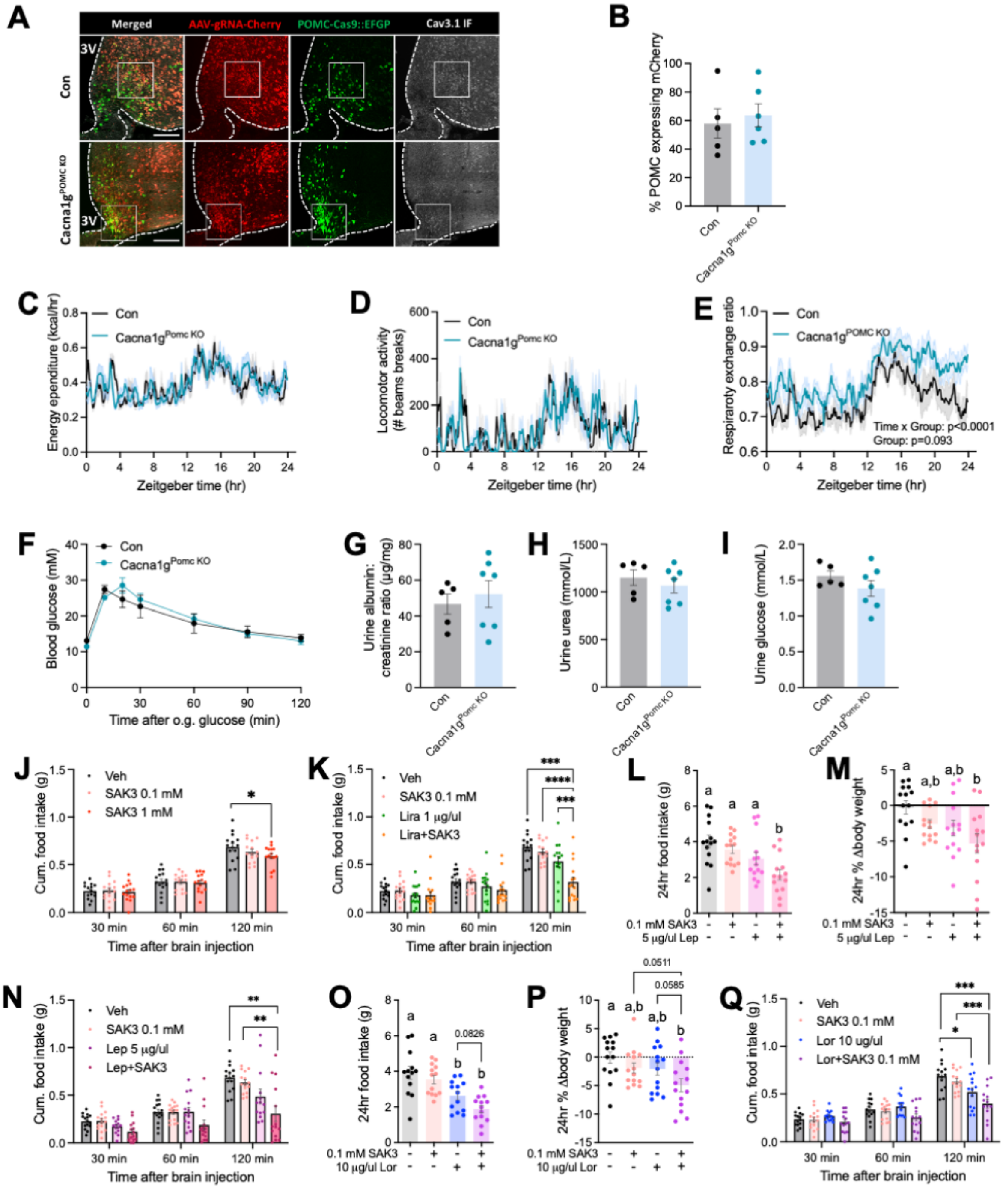

**Figure S6: Additional data related to Figure 5-6**

**(A-B)** Confocal **i**mmunofluorescence (IF) microscopy analysis of the CRISPR knockout efficiency of Cav3.1 in Cacna1g$^{POMC\ KO}$ mediobasal hypothalamus with a C-terminal specific Cav3.1 antibody validated in Fig. S3F. (**A**) Zoom-out images of Fig. 5B. Scale bars: 200 µm. (**B**) Quantification of hypothalamic POMC neurons transduced with AAVs. Con: n = 5; Cacna1g$^{POMC\ KO}$: n = 6. **(C-E)** Metabolic parameters of Cacna1g$^{POMC\ KO}$ mice on P45 diet. (**C**) 24 hr profile of energy expenditure, (**C**) 24 hr profiles of locomotor activity and (**D**) respiratory exchange ratio (**E**) during indirect calorimetry measurement. Con: n = 5; Cacna1g$^{POMC\ KO}$: n = 7. **(F)** Oral glucose tolerance test of Con and Cacna1g$^{MBH\ KO}$ mice on P14 diet. Con: n = 5; Cacna1g$^{POMC\ KO}$: n = 7. **(G-I)** Urine analyses of Con and Cacna1g$^{POMC\ KO}$

mice fed on P45 diets. Urinary albumin/creatinine ACR ratio (**G**), urea (**H**) and glucose (**I**). Con: n = 5; Cacna1g$^{POMC\ KO}$: n = 7. **(J)** Acute feeding responses after MBH injection of Veh, 0.1 mM and 1 mM $Ca_v3.1$ activator SAK3. n = 14 per group. **(K)** Acute feeding responses after MBH injection of Veh, 0.1 mM SAK3, 1 µg/µl liraglutide (Lira) and SAK3+Lira co-treatment. n = 14 per group. **(L-N)** 24 hr feeding responses (**L**), 24 hr % body weight changes (**M**) and acute feeding responses (**N**) after MBH injection of Veh, 0.1 mM SAK3, 5ug/µl leptin (Lep) and SAK3+Lep co-treatment. n = 14 per group. **(O-Q)** 24 hr feeding responses (**O**), 24 hr % body weight changes (**P**) and acute feeding responses (**Q**) after MBH injection of Veh, 0.1 mM SAK3, 10ug/µl lorcaserin (Lor) and SAK3+Lor co-treatment. n = 14 per group. Groups denoted with different letter in (L, M, O, P) indicate significant difference (p<0.05). *p<0.05, **p<0.01, ***p<0.001, ****p<0.0001. Values are reported as mean ± SEM.
